# Synapse-specific diversity of distinct postsynaptic GluN2 subtypes defines transmission strength in spinal lamina I

**DOI:** 10.1101/2023.03.02.530864

**Authors:** Graham. M. Pitcher, Livia Garzia, A.S. Morrissy, Michael D. Taylor, Michael. W. Salter

**Affiliations:** Program in Neurosciences & Mental Health, The Hospital for Sick Children, Toronto, ON, Canada; Department of Physiology, University of Toronto, University of Calgary, Canada; Department of Surgery, Faculty of Medicine, McGill University, and Cancer Research Program, The Research Institute of the McGill University Health Centre, University of Calgary, Canada; Department of Biochemistry & Molecular Biology, Cumming School of Medicine, University of Calgary, Canada; Brain Tumor Program, Texas Medical Centre, Houston, TX, U.S.A.

## Abstract

The unitary postsynaptic response to presynaptic quantal glutamate release is the fundamental basis of excitatory information transfer between neurons. The view, however, of individual glutamatergic synaptic connections in a population as homogenous, fixed-strength units of neural communication is becoming increasingly scrutinized. Here, we used minimal stimulation of individual glutamatergic afferent axons to evoke single synapse resolution postsynaptic responses from central sensory lamina I neurons in an *ex vivo* adult rat spinal slice preparation. We detected unitary events exhibiting a NMDA receptor component with distinct kinetic properties across synapses conferred by specific GluN2 subunit composition, indicative of GluN2 subtype-based postsynaptic heterogeneity. GluN2A, 2A and 2B, or 2B and 2D synaptic predominance functioned on distinct lamina I neuron types to narrowly, intermediately, or widely tune, respectively, the duration of evoked unitary depolarization events from resting membrane potential, which enabled individual synapses to grade differentially depolarizing steps during temporally-patterned afferent input. Our results lead to a model wherein a core locus of proteomic complexity prevails at this central glutamatergic sensory synapse that involves distinct GluN2 subtype configurations. These findings have major implications for subthreshold integrative capacity and transmission strength in spinal lamina I and other CNS regions.

## INTRODUCTION

The unitary postsynaptic response to quantal glutamate release is the fundamental basis of excitatory information transfer in the central nervous system (CNS). The prevailing assumption that glutamatergic synaptic connections in a given neural network or population function as homogenous, equal strength relay units of excitatory information is beginning to face critical re-evaluation^1^. The core components of a typical glutamatergic synapse include glutamate-gated ionotropic receptors (iGluRs) consisting primarily of α-amino-3-hydroxy-5-methyl-4-isoxazole propionate (AMPA) and N-methyl-D-aspartate glutamate receptor-channel (NMDAR) complexes, along with adhesion and signaling molecules ^2–4^. However, there is increased scrutiny directed towards a more comprehensive understanding of postsynaptic molecular composition and assembly – do a multitude of postsynaptic receptor-channel and protein constituents occur as a generic mix across synapses in a population, or is there a molecular codification to their nanopositioning and function that tunes transmission strength at individual glutamatergic synapses?

Recent advances in single-cell RNA sequencing and proteomics in the rodent brain^2, 4–6^ and spinal cord^7, 8^ have identified considerable diversity in synaptic RNA and protein content. At glutamatergic synapses in particular, molecular complexity is reflected extensively in the postsynaptic density (PSD) portion^2, 4, 8–10^ where AMPA and NMDA receptors are optimally situated for synaptic function. Though the NMDAR multiprotein structure itself is composed exclusively of ion channel subunits consisting of a heterotetramer core comprised of two obligate GluN1 subunits and two GluN2 subunits ^11^, NMDARs contained within ‘NMDAR supercomplexes’ ^3, 12^ are bound to a myriad of adhesion and signalling binding partners ^3, 12, 13^. Notably, dissimilar GluN2 subunits of which four types exist (2A-D) ^11^ are subjected to differing interactions with distinct PSD proteins ^14–19^ including for example PSD-95 ^20–23^ indicative of GluN2 subtype regulation by distinct PSD binding partners. Although there is compelling evidence that NMDARs congregate in a cluster primarily at the center of the PSD ^24–26^, recent progress *in vitro* using single-molecule super-resolution imaging in cultured hippocampal neurons has now identified more precise spatial organization of distinct GluN2 subtypes in defined nanodomains in the PSD with the localization of each determined individually by specific PSD binding partners within the NMDAR supercomplex ^27^. The known extensive complexity of PSD protein constituents ^2, 3, 5^ and the regulatory function they impose on GluN2 subtype nanoscale localization *in vitro* ^27^ therefore open the potential for multiple scenarios of diverse GluN2 subtype(s) organization within the PSD compartment. However, our current understanding of GluN2 subtype diversity and distinct PSD compartmentalization at the resolution of individual synapses in higher-order mammalian synaptic circuits or populations remains elusive. Moreover, though GluN2 subunit composition within the NMDAR complex is known to determine several biophysical receptor-channel properties that mediate current flow ^28, 29^, the functional significance that distinct GluN2 subtype localization within the PSD imposes on unitary integrative capacity and transmission strength at individual synapses remains unexplored. Limited consideration may be dedicated to these issues likely because of the technically challenging nature of investigating NMDARs, their properties, and function at single-synapse resolution in *in vivo* or *ex vivo* CNS preparations.

We set out here to understand GluN2 subtype organization and function at the resolution of individual mature mammalian glutamatergic synaptic connections using an electrophysiological approach. We developed a minimal stimulation protocol in an *ex vivo* adult rat spinal cord slice preparation in which we evoked unitary excitatory postsynaptic currents (μEPSCs) and potentials (μEPSPs) at individual glutamatergic monosynaptic connections ^30–33^ between primary sensory afferents in the dorsal rootlet and central neurons in the lamina I nociceptive subfield of the spinal dorsal horn, a major CNS somatosensory region. μEPSCs exhibited distinctive NMDAR kinetic properties at this principal sensory synapse providing evidence supporting a high degree of diversity of single or multiple GluN2 subtype postsynaptic configurations at individual glutamatergic synaptic connections. μEPSP recordings further identified distinct GluN2 subtype configurations as a fundamental single-synapse resolution coding mechanism. Our results support a model wherein the mature mammalian primary afferent-lamina I neuron glutamatergic synapse is a core locus of distinct GluN2 subtype-based proteome diversity and nanoscale organization that directly influences fundamental aspects of subthreshold transmission efficacy including unitary resolution synaptic computation capability and strength, and ultimately how somatosensory information is processed. In other CNS regions diverse postsynaptic GluN2 subunit composition may constitute distinct integrative signatures at glutamatergic synapses that bear major implications in physiological transmission strength and information transfer.

## RESULTS

### Unitary primary afferent-lamina I synaptic responses

To investigate NMDAR GluN2 subtypes at mature, lamina I synapses, we made whole-cell voltage-clamp recordings from visually identified lamina I neurons in acute *ex vivo* dorsal root-attached transverse spinal cord slices from adult rats. We elicited primary-afferent mediated post-synaptic responses in lamina I neurons by delivering single electrical stimuli to the attached dorsal rootlet. Stimulation of primary afferents synaptically connected to lamina I neurons evoked excitatory post-synaptic currents (EPSCs) (Fig. 1a). When the membrane potential was held at hyperpolarized potentials, we observed a rapid-onset, rapidly decaying EPSC response. Holding the neuron at positive membrane potentials revealed a slowly decaying component of the EPSC, in addition to the rapid-onset peak. The peak of the rapid onset component exhibited a linear current-voltage relationship, whereas the slowly-decaying EPSC component showed dramatic inward rectification (Fig. 1a). The rapid-onset component of the EPSC was blocked by the AMPAR antagonist CNQX and the slow component was blocked by the NMDAR competitive antagonist D-amino-phosphonovaleric acid (D-APV, 100 μM; not illustrated). Thus, adult lamina I dorsal horn neurons exhibit primary afferent-evoked EPSCs with kinetic and pharmacological properties of both AMPA and NMDA receptors.

**Figure 1.**
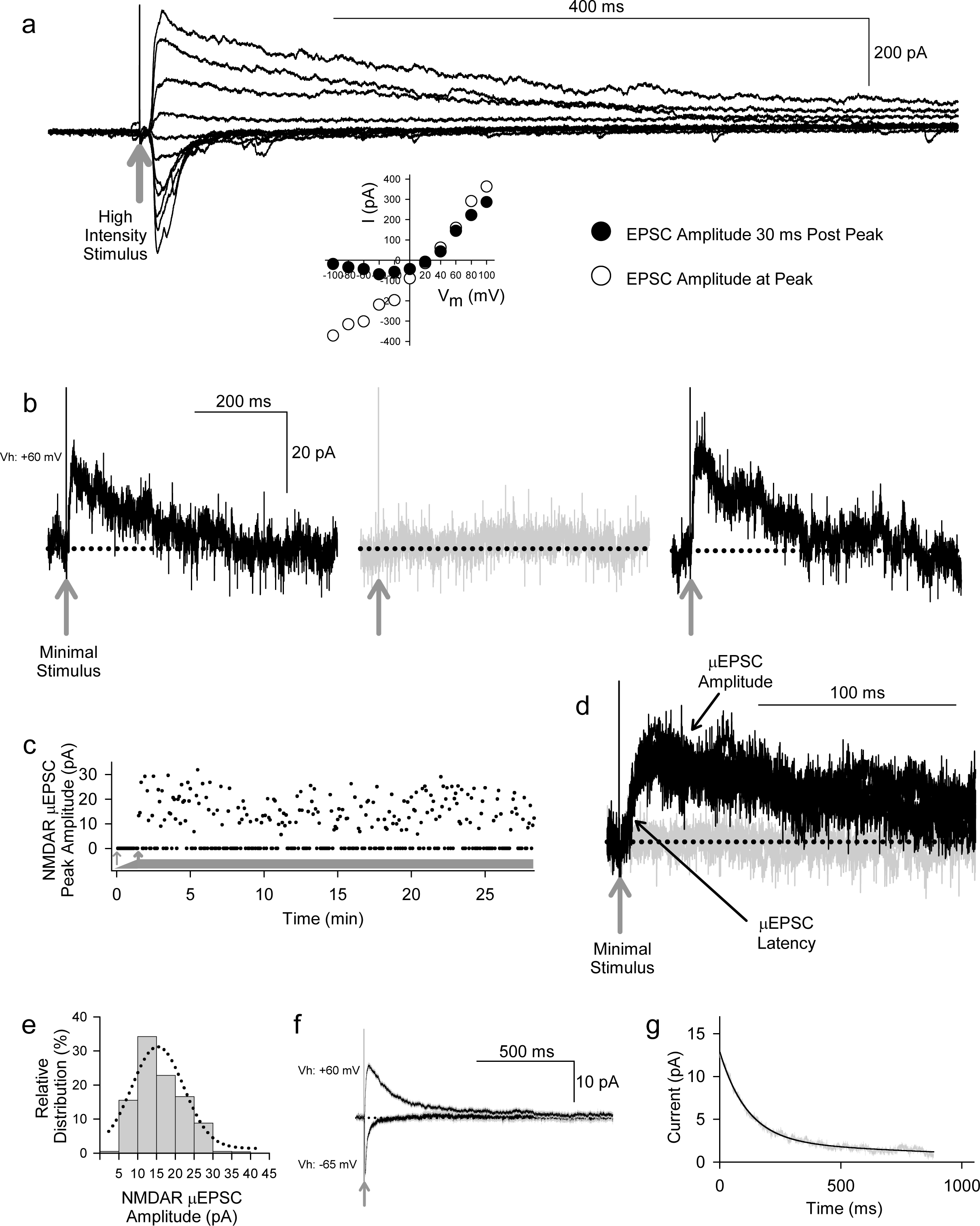
Primary afferent-evoked μEPSCs in whole cell voltage-clamp recordings from lamina I neurons in acute *ex vivo* dorsal root-attached transverse spinal cord slices from adult rats exhibited kinetic properties of both AMPA and NMDARs. (a) Delivering single electrical stimuli to the attached rootlet that activated multiple primary afferents evoked EPSCs in lamina I neurons. Stimulation at hyperpolarized membrane potentials elicited rapid onset, rapidly decaying EPSC responses, whereas stimulation at depolarizing membrane potentials evoked a slowly decaying component, in addition to the rapid-onset peak. Peak of the rapid onset component exhibited a linear current-voltage relationship (inset below, white symbols) consistent with kinetic properties of AMPARs, while the slowly decaying EPSC component exhibited inward rectification (inset below, black symbols) consistent with current-voltage (I-V) relationship of NMDARs. (b) Recordings of unitary-evoked EPSCs from a representative adult rat spinal lamina I neuron elicited by minimal stimulation of individual axons in the attached dorsal rootlet. Sample traces showing individual consecutive all-or-none μEPSCs held at +60 mV evoked by minimal stimulation. The black traces show successful synaptic μEPSCs in response to minimal sensory afferent stimulation (gray arrow indicates stimulus artefact) while the gray trace shows a synaptic failure. (c) Scatter plot of successful synaptic μEPSC responses and failures over time (thick gray bar) evoked by minimal stimulation (0.2 Hz). The thin gray arrow indicates subthreshold stimulation intensity while the thick gray arrow indicates threshold at which μEPSCs can be evoked. (d) Several successful synaptic μEPSCs (black traces) from the same recording in (b) showing identical amplitude and latency (black arrows) consistent with a monosynaptic connection (gray arrow indicates stimulus artefact). The gray trace indicates a synaptic failure. (e) NMDAR amplitude histogram showing relative distribution for successful μEPSCs, fit with a single with a single Gaussian function. (f) Averaged traces (± SEM) showing successful μEPSCs held at +60 mV and successful μEPSCs recorded at -65 mV. (g) Single exponential fitting of the decay component of the averaged μEPSC held at +60 mV.

To examine individual primary-afferent evoked EPSCs of lamina I neurons in the dorsal root-attached spinal cord slice preparation we used a minimal stimulation protocol ^34^ to evoke unitary, monosynaptic EPSCs (μEPSCs) in the lamina I neuron recorded (n=224 cells). The criteria used to establish monosynaptic μEPSC responses were: a distinct electrical stimulation threshold at which responses were all-or-none (ie. success or failure; Fig. 1b-c), a consistent latency throughout the duration of each recording (Fig. 1d), and the amplitude of the NMDAR component was normally distributed (Fig. 1e). We confirmed that responses evoked by the minimal stimulation were *bona fide* unitary responses because increasing stimulus intensity did not increase μEPSC amplitude until the intensity was sufficient to evoke a second level of all-or-none evoked responses (Supplementary Fig. 1c). Moreover, bath-applying tetrodotoxin (TTX) to block voltage-gated Na^+^ channels eliminated the μEPSCs (Supplementary Fig. 1a-b), indicating that unitary synaptic responses were driven by action potentials evoked by direct stimulation of a presynaptic axon in the dorsal rootlet rather than by direct focal stimulation of the lamina I neuron.

To determine whether or not the response failures with minimal stimulation were due to failure of the stimulating electrode to generate an action potential in the primary afferent axon in the dorsal rootlet eliciting the μEPSC, we used paired-pulse minimal stimulation experiments (Supplementary Fig. 1d-e). We found that the average amplitude of the μEPSCs to the second stimulus was greater than that of the first when the first stimulus also evoked a μEPSC (Supplementary Fig. 1d-e, left panels), consistent with paired-pulse facilitation of transmitter release. Moreover, the average amplitude of the μEPSCs to the second stimulus that followed a failure of response to the first stimulus was identical to, if not greater than, the average of the responses to the second stimulus that followed first stimulus non-failures (Supplementary Fig. 1d-e, right panels). These findings are not consistent with failure of the first stimulus to generate an action potential, in which case the amplitude of the μEPSC to the second stimulus would have been predicted to be the same as that of the first stimulus non-failures. Rather, our findings show that despite the failure of post-synaptic response to the first stimulus an action potential must have been triggered in the axon connected to the lamina I neuron, and must have reached the presynaptic terminal to enhance quantal release of glutamate presynaptically in response to the second stimulus. Therefore, we conclude that the minimal stimulation technique used here reliably evoked a monosynaptic, excitatory post-synaptic response from a single primary afferent connected to each lamina I neuron studied.

When initially searching for a unitary synaptic input in μEPSC recordings we held the membrane potential at -65 mV (n=172 cells). In all of these recordings when the membrane potential was subsequently held at +60 mV we observed a component of the decay phase that was slower than that when the neuron was held at -65 mV, which we interpret as indicating that there were both AMPAR- and NMDAR-components to the μEPSCs. In the remainder of recordings we searched with the membrane potential held at +60 mV to look for NMDAR-only synapses.

However, in all cases we found a rapid inward current when the neuron was subsequently held at - 65 mV. Thus, we conclude that all single, primary-afferent-to-lamina I neuron synapses we recorded contained both AMPAR and NMDAR components.

### Three distinct **μ**EPSC types evoked across individual adult primary afferent-lamina I neuron synapses

μEPSC average amplitude and decay were found to be stable within each single synapse recording (Vh: +60 mV) throughout the duration of the recording period (e.g. Fig. 1b-d,f-g). However, we observed considerable heterogeneity in the μEPSC decay *between* single-synapse recordings (Fig. 2a). To gain insight into the diversity in μEPSCs between recordings we performed principal component analysis (PCA) followed by hierarchical clustering (HC). For the PCA we examined several parameters including charge transfer, half-width, decay constant, peak amplitude, rise time, neuronal input resistance, and synaptic failure rate. Dimensionality reduction of the parameters with PCA (Fig. 2b-c and Supplemental Fig. 2), showed that the first three components capture 76.1% of the data variability. Based on this, μEPSC decay, charge transfer and half-width were retained as electrophysiological parameters that significantly contributed to the first three PCA components (Figure 2c) and, therefore, used to build a classification of individual synaptic responses. To determine whether primary afferent-lamina I synapses are comprised of distinct types, we used μEPSC decay, charge transfer and half-width for the HC (Fig. 2d). Gap-statistic analysis of the clustering (Fig.2d inset top right) identified three as the optimal number of clusters to represent the dataset. We labelled these three groups: fast (n=80, 36% of recorded neurons), intermediate (n=114, 51% of recorded neurons), and slow (n=30, 13% of recorded neurons) based on decay time constant of their μEPSCs (Fig. 2e). Individual representative fast, intermediate, and slow μEPSCs are shown in Supplementary Figure 3. From these representative μEPSCs, latency, amplitude, and decay can be seen to be stable throughout each recording (Supplementary Fig. 4). Mean μEPSC traces for each of the three unitary synaptic response types determined by clustering are shown in Figure 3a-c. Across these three groups the following electrophysiological parameters were not different: failure rate (fast 56±2%, intermediate 52±2%, and slow 51±3%), cell input resistance (fast 363±32 MOhm, intermediate 334±19 MOhm, and slow 367±34 MOhm) as well as μEPSC rise time (fast 2.9±0.2 ms, intermediate 3.1±0.2 ms, and slow 3.4±0.3 ms). Therefore, we conclude that primary afferent-lamina I μEPSCs at the holding potential of +60 mV can be classified into three major groups according to decay constant, half-width, and charge transfer (Fig. 3d).

**Figure 2.**
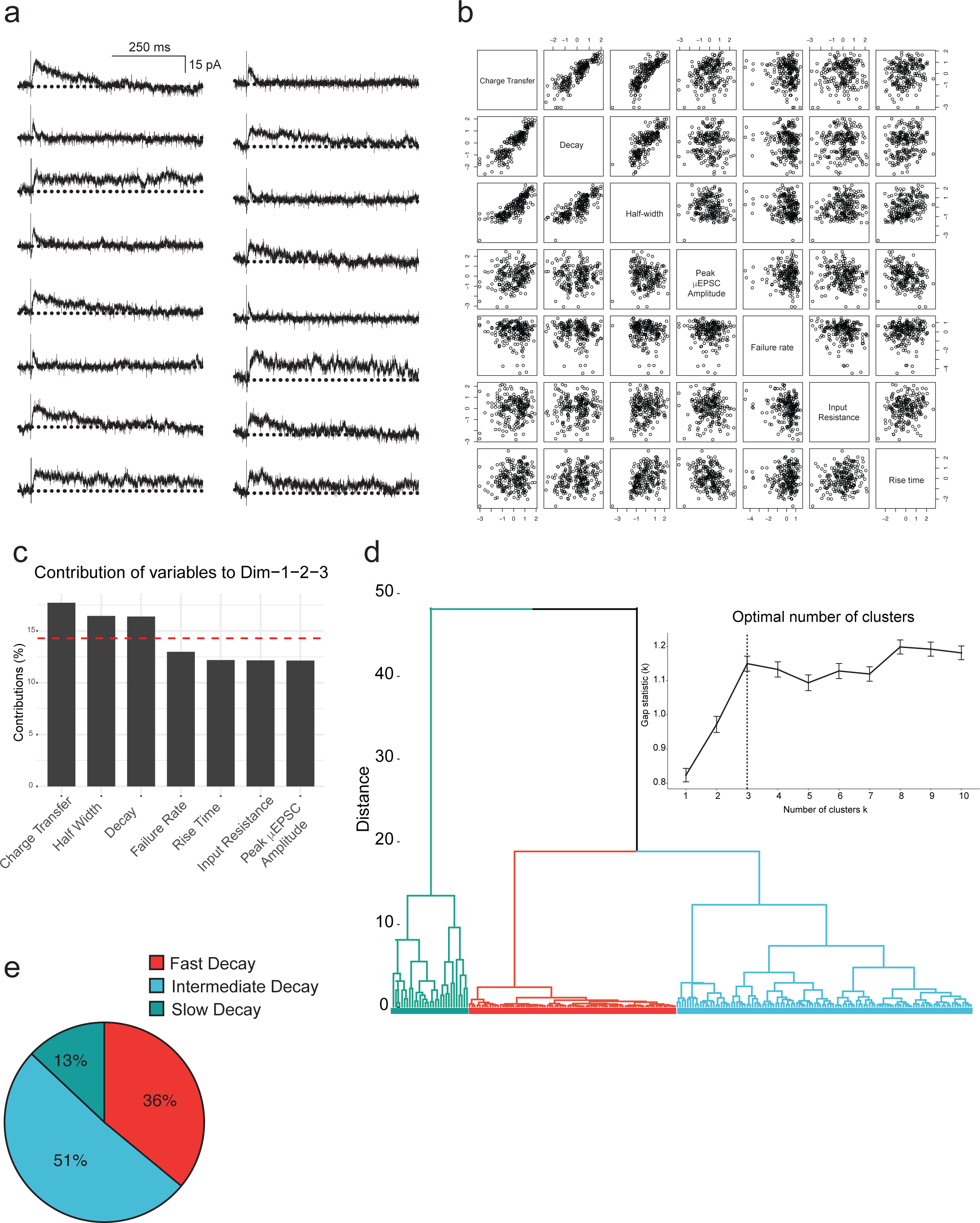
Principal component and hierarchical cluster analysis of the electrophysiological parameters of the μEPSCs recorded (Vh: +60 mV) (a) Representative μEPSCs recorded at +60 mV illustrating the diversity of the kinetic properties. (b) Correlation plots of the variables in this study incorporated in the PCA analysis of the 224 independent recordings. (c) Histogram representing the contribution of variables to the three first dimensions, representing 76% of the data variance. The red dashed line on the graph indicates the expected average contribution if the contribution of the variables were uniform. (d) Heirchical clustering and gap-statistic analysis (inset) of the three first components identify three clusters of responses. © The pie chart shows relative (percent) prevalence of distinct mEPSC types in the full dataset.

**Figure 3.**
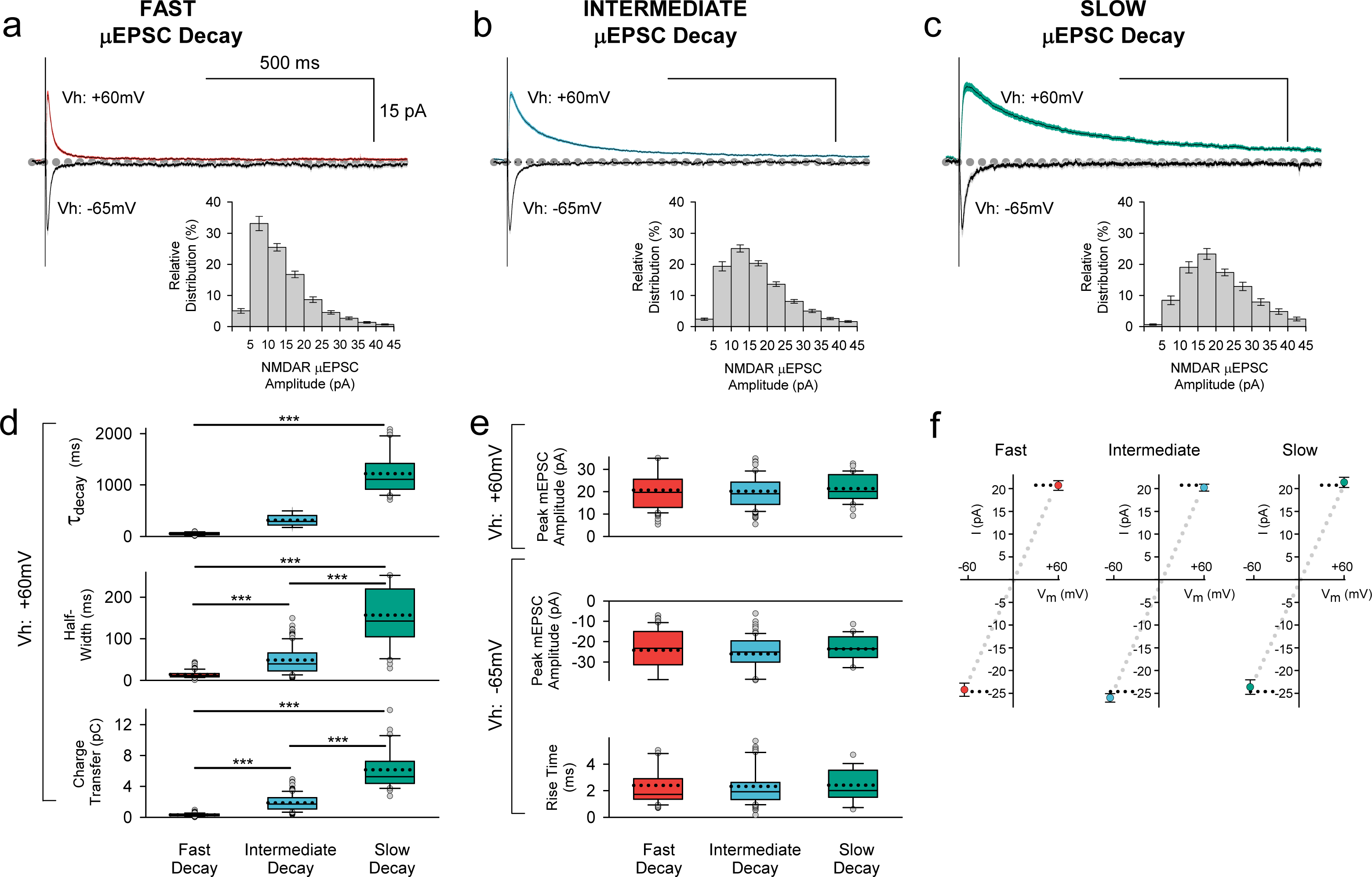
Mean traces for each of the three unitary synaptic response types determined by cluster analysis. (a) Top: Averaged traces (± SEM) showing successful fast decay μEPSCs (n = 80) at +60 mV (red) and -65 mV (gray). Insert at bottom: Mean NMDAR amplitude histogram showing relative distribution for successful fast decay μEPSCs. (b) Top: Averaged traces (± SEM) showing successful intermediate decay μEPSCs (n = 114) at +60 mV (blue) and -65 mV (gray). Insert at bottom: Mean NMDAR amplitude histogram showing relative distribution for successful intermediate decay μEPSCs. (c) Top: Averaged traces (±SEM) showing successful slow decay μEPSCs (n = 30) at +60 mV (green) and -65 mV (gray). Insert at bottom: Mean NMDAR amplitude histogram showing relative distribution for successful slow decay μEPSCs. (d) Summary of NMDAR-mediated μEPSC (Vh: +60 mV) decay (top), half-width (middle), and charge transfer (bottom) for fast decay (red), intermediate (blue) and slow decay (green) μEPSCs (p < 0.001, unpaired, two-tailed t-test). Dotted lines in box plots indicates mean. (e) Top: Summary of AMPAR-mediated μEPSC peak amplitude (Vh: +60 mV) for fast, intermediate, and slow unitary responses. Bottom: Summary of AMPAR-mediated μEPSC peak amplitude and rise time (Vh: -60 mV) for fast, intermediate, and slow unitary responses. (f) Mean (±SEM) current-voltage (I-V) relationship for μEPSC peak amplitudes held at +60 mV and -65 mV for fast, intermediate, and slow groups.

Across these three groups, the μEPSCs at the holding potential of -65 mV did not differ in peak amplitudes (fast -24±1.4 pA, intermediate -26±0.9 pA, and slow -24±1.6 pA; p > 0.05) or in rise times (fast 2.4±0.2 ms, intermediate 2.3±0.2 ms, and slow 2.4±0.3 ms; p > 0.05; Fig. 3a-c below; Fig. 3e). Moreover, the peak amplitudes of μEPSCs held at -65 mV exhibited linear current-voltage relationships (Fig. 3f). Thus, the AMPAR components of the μEPSCs were indistinguishable across the three groups.

### Distinct GluN2 subtypes mediate the decay kinetics of the three distinct unitary synaptic responses

As the NMDAR component but not the AMPAR component of the uEPSCs fell into three groups we investigated the basis for the differences in the NMDAR component. Of the three kinetic parameters that account for variance within the population of μEPSCs, half-width and charge transfer (calculated as area under the curve) occur as a passive function of the time course of the μEPSC decay (Fig. 3). Thus, we inferred from the readily discernible correlation of charge transfer and half-width with decay that the most critical parameter governing the functional importance of each unitary response is time course of the decay itself. We therefore focused our attention on elucidating the mechanism(s) responsible for different μEPSC decay kinetics.

Notably, given that the three distinct μEPSC decay constants determined from our cluster analysis (52±3 ms for ‘fast’ decay; 315±11 ms for ‘intermediate’ decay; 1219±69 ms for ‘slow’ decay at Vh: +60mV; p < 0.001; Fig. 3d) correspond well with values from known NMDAR GluN2 subtypes including specifically GluN1/2A/2A, 2B/2B, and 2D/2D, respectively ^34, 35^, we hypothesized that the dissimilar, yet D-APV-sensitive (Supplementary Fig. 5), decay components of unitary synaptic responses represent distinct deactivation kinetics governed by differences in intrinsic properties of distinct GluN2 subtype-containing NMDARs at these synapses.

To test this hypothesis, we rigorously investigated the pharmacological sensitivity of the three distinct response types to known selective NMDAR GluN2 subtype inhibitors. We tested first involvement of GluN2D/2D-containing NMDARs. In a subset of voltage-clamp recordings examining representative ‘fast’, ‘intermediate’, and ‘slow’ μEPSC responses (Vh: +60 mV), we administered to aCSF the GluN2D/2D selective antagonist, DQP-1105 (10 μM), which is selective for GluN2D/2D diheteromeric NMDARs^36^ which we refer to here as GluN2D. Fast and intermediate unitary responses were not affected by DQP-1105 administration (Supplementary Fig. 6b-d,f,g), whereas the duration of ‘slow’ unitary kinetic responses was reduced by DQP-1105 (Supplementary Fig. 6a,d,e). Charge transfer of the NMDAR component measured with DQP-1105 present in aCSF was reduced significantly compared to charge transfer during baseline μEPSC responses before DQP-1105 administration (p < 0.05; Supplementary Fig. 6d-e). Taken together, these results indicate that the particularly prolonged decay of the ‘slow’ μEPSC response was mediated by GluN2D-containing NMDARs.

We investigated next possible involvement of GluN2B/2B-containing NMDARs by bath administering the GluN2B/2B selective antagonist, Ro25-6981 (1 μM) which is selective for GluN2B/2B diheteromeric NMDARs ^37^ which we refer to here as GluN2B. Ro25-6981 was without effect on ‘fast’ decay μEPSC responses (Supplementary Fig. 7c-d,g) whereas the duration of the NMDAR component of ‘intermediate’ decay μEPSC responses was reduced; the NMDAR charge transfer with Ro25-6981 present was decreased significantly (p < 0.01) compared to charge transfer during baseline μEPSC responses prior to Ro25-6981 administration (Supplementary Fig. 7b,d,f). For ‘slow’ decay responses the intermediate portion of the NMDAR component was affected only moderately by Ro25-6981 while the late NMDAR component of the μEPSC decay was unaffected (Supplementary Fig. 7a); charge transfer of the NMDAR component was reduced by Ro25-6981 compared to baseline μEPSC responses (p < 0.001; Supplementary Fig. 7d-e). We therefore conclude that both ‘intermediate’ and ‘slow’ synapse types are comprised, at least in part, of GluN2B-containing NMDARs.

Having shown that the NMDAR component of ‘fast’ μEPSC responses was resistant to both Ro25-6981 and DQP-1105 (Supplementary Figs. 6-7), we reasoned that ‘fast’ unitary primary afferent-lamina I synaptic responses are mediated by NMDARs other than GluN2B- or GluN2D-containing subtypes. We considered, therefore, the possibility that GluN2A/2A- containing diheteromeric NMDARs which are known to exhibit decay kinetics faster than those of the other GluN2 subunits ^34^ may underlie this NMDAR-mediated synaptic response. To investigate potential function of GluN2A/2A-containing NMDARs (which we refer to here as GluN2A) at these synapses we administered the selective antagonist, TCN-201 (3 μM) ^38^. We tested TCN-201 on representative ‘fast’ decay μEPSC responses and found that the μEPSC duration was reduced (Supplementary Fig. 8a-c) to the extent that the entire charge transfer mediated by NMDAR function was abolished (p < 0.01; Supplementary Fig. 8d-e). We conclude that GluN2A-containing NMDARs comprise the ‘fast’ synapse type.

The depressive effect of Ro25-6981 on slow decay μEPSCs (Supplementary Fig. 7a,d-e) suggests function of GluN2B-containing in addition to GluN2D-containing NMDARs at ‘slow’ decay synapses. In addition, that the residual ‘DQP-1105-insensitive’ NMDAR component of ‘slow’ decay μEPSCs showing sensitivity to DQP-1105 is comparable to ‘intermediate’ μEPSC responses (Supplementary Fig. 6a,d-f) further provides evidence for GluN2B subtype NMDARs in addition to GluN2D-containing NMDARs at ‘slow’ decay synapses. Furthermore, though we did not examine specifically TCN-201 on ‘intermediate’ and ‘slow’ decay μEPSC responses, we are not ruling out a contribution of GluN2A-containing NMDAR function at these synapses. For example, the τ_(fast)_ for ‘intermediate’ decay μEPSCs (45±4 ms) is comparable (p > 0.05) to the τ value for ‘fast’ (GluN2A-mediated) μEPSC decay kinetics (52±3 ms; Fig. 3d). It is also evident from ‘intermediate’ decay μEPSCs which showed sensitivity to Ro25-6981 that the remaining ‘Ro25-6981-insensitive’ NMDAR component is comparable to the ‘fast’ decay μEPSC responses (Supplementray Fig. 7c-d,f-g). Based on these observations, we are not excluding the possibility that ‘intermediate’ and possibly ‘slow’ synapses may also contain GluN2A-containing NMDARs.

### Differential Mg^2+^ block of fast, intermediate, and slow **μ**EPSCs

GluN2 subunit differences in the kinetics of Mg^2+^ unblock contribute to NMDAR subtype function in current flow during membrane depolarization at glutamatergic synapses ^28^. Given the distinct GluN2 kinetic properties determined from the three μEPSC types recorded at the holding potential of +60 mV (Fig. 3a-d), we reasoned that μEPSCs recorded from the same experiments but at the negative holding potential of -65 mV (Fig. 3a-c bottom) would also reflect differences in biophysical properties indicative of distinct GluN2 subtypes. Consistent with this prediction, the late component of negatively held μEPSCs exhibited significant differences in half-width (p < 0.01) and charge transfer (p < 0.05, 0.01, 0.001) across the three groups (Supplementary Fig. 9a) that may be attributed to differences in sensitivity of GluN2 subtypes to Mg^2+^ block ^28, 39^. By determining the Mg^2+^-induced percent inhibition of the amplitude of the NMDAR component of μEPSCs recorded at the holding potential of -65 mV in each group (Supplementary Fig. 9b) we found that the inhibition of the NMDAR component of μEPSCs from the ‘fast’ cluster was similar to that of μEPSCs from the ‘intermediate’ cluster, whereas the Mg^2+^-induced percent inhibition of the NMDAR component of μEPSCs from the ‘slow’ cluster was substantially less (Supplementary Fig. 9b) ^28, 39^. We conclude that the differences in Mg^2+^ block of μEPSC NMDAR components at primary afferent-lamina I glutamatergic synapses further confirm the distinctiveness of GluN2-containing NMDAR subtype expression and function we propose to occur at these synapses.

### No evidence for presynaptic NMDAR function

We reasoned that if NMDARs functioned presynaptically to influence synaptic transmission, we would observe a change in unitary synaptic failure rate. We found that failure rate remained stable in recordings in which D-APV, TCN-201, Ro25-6981, or DQP-1105 was tested (Supplementary Fig. 10a-c). In a representative experiment, D-APV is shown to inhibit the NMDAR component of the μEPSC (mean±SEM) while exhibiting no effect on failure rate (Supplementary Fig. 10a) consistent with a postsynaptic function of NMDARs at primary afferent-lamina I neuron synapses.

### **μ**EPSP temporal summation at individual synapses is NMDAR-dependent

The diversity of GluN2-containing NMDARs at individual lamina I synapses and their differences in Mg^2+^ block prompted us to investigate next function of GluN2 subtype diversity at individual primary afferent-lamina I synapses. We carried out minimal stimulation in voltage-clamp first to establish *bona fide* μEPSC responses (Vh: -65 mV) after which, within the same recording, we switched to current-clamp mode to monitor membrane potential (μEPSP response using a K^+^- based intracellular solution). In this dual mode recording protocol we used a brief train composed of 5 stimuli delivered every 200 ms (5 Hz frequency which mimics low repetitive firing pattern of nociceptors ^40^) at 5 s intervals (0.2 Hz) in voltage clamp (Supplementary Fig. 11a-b) and then current clamp (Supplementary Fig. 11c-d) modes for a given lamina I recording.

Continuous sweep recordings of μEPSCs during the 5 Hz minimal stimulus train show that mean (±SEM) peak amplitude, half-width, and rise time remained stable (p > 0.05; Supplementary Fig. 11 a-b). Though an increase in μEPSC mean (±SEM) charge transfer was observed over successive stimuli (p < 0.01), pure μEPSC responses (without failures) at each stimulus throughout the 5 Hz train remained stable; peak amplitude, charge transfer, half-width, and rise time for the five μEPSCs in the 5 Hz train were not significantly different (p > 0.05; Supplementary Fig. 12a-b). A decrease in failure rate was found between the first and fifth pure μEPSCs (successes only) in the 5 Hz train (p < 0.05; Supplementary Fig. 12b). Though indicative of increased presynaptic glutamate release probability through the progression of the 5 Hz train, we do not consider this to have influenced postsynaptic mechanisms during the train because each pure μEPSC response (without failures) remained constant.

Continuous sweeps recorded in current clamp mode during the 5 Hz train showed that membrane potential depolarized progressively with each stimulus, peaking after the fifth stimulus, a response we refer to as a ‘μEPSP burst’ (Supplementary Fig. 11c-d). Pure μEPSP responses (successes only) shown at each stimulus (mean±SEM) during the 5 Hz train exhibited a progressive increase in peak amplitude, area, half-width, decay, and prestimulus membrane potential (p < 0.001; Supplementary Fig. 13a-b), while rise time across the five μEPSPs in the train was not different (p > 0.05; Supplementary Fig. 13a-b). Notably, the failure rates for the first and fifth μEPSPs were indistinguishable from those for the corresponding first and fifth μEPSCs recorded in voltage clamp (p > 0.05; Supplementary Fig. 12b) which indicate that the transition in recording mode (from voltage clamp) was inconsequential to mechanisms underlying unitary synaptic transmission as we were able to evoke reliable μEPSPs that summate with a failure rate that was indiscernible from that during μEPSC recordings.

In a representative example μEPSP burst, D-APV was without effect on the rising phase or the peak amplitude of the first μEPSP in the 5 Hz train but reduced μEPSP decay and abolished temporal summation (Supplementary Fig. 14a). Similarly, the NMDAR channel blocker, MK-801, reversed the summated component of the μEPSP burst (Supplementary Fig. 14b). Taken together, we conclude that temporally-patterned μEPSPs during the 5 Hz stimulus train exhibit summation and that μEPSP decay and summation are dependent on NMDAR function.

### Distinct μEPSP kinetics across individual primary afferent-lamina I neuron synapses

Having determined that that temporal summation of μEPSPs at this synapse is attributable to NMDAR function (Supplementary Fig. 14), we reasoned that GluN2 subtype synaptic distinctiveness (Figs. 2-3; Supplementary Figs 2-9) may underlie possible heterogeneity in our μEPSP burst dataset. In one approach to elucidate potential diversity in μEPSP burst properties, we grouped μEPSP bursts based on lamina I neuron type (Fig. 4a-c). In each current-clamp recording, in which we had acquired also membrane potential in response to depolarizing current injection (20 pA for 1 s), we identified in lamina I neurons distinct firing properties in response to current injection. This enabled us a subclassification of our μEPSP dataset based on neuron spiking properties including single/non-spiking, delayed/phasic spiking, or tonic spiking activity (Fig. 4a-c), established features of lamina I neurons ^41–43^. We examined kinetic properties from the first μEPSP in the burst as it was the μEPSP in the stimulus train without summation, and from the last (fifth) μEPSP in the burst as it exhibited the greatest degree of summation (Supplementary Fig. 13). Within lamina I neuron type, pure μEPSPs (successes only, mean±SEM) exhibited an increase in area, half-width, decay, and prestimulus membrane potential of the fifth μEPSP vs the first μEPSP (p < 0.05, p < 0.01, p < 0.001, Fig. 4e-h). Between lamina I neuron type, pure first and fifth μEPSPs exhibited differences in area, half-width, and decay (p < 0.05, p < 0.01, p < 0.001; Fig. 4e-g). Pure fifth μEPSPs between neuron type also exhibited significant differences in peak amplitude and prestimulus membrane potential (p < 0.05, p < 0.01, p < 0.001; Fig. 4d,h).

**Figure 4.**
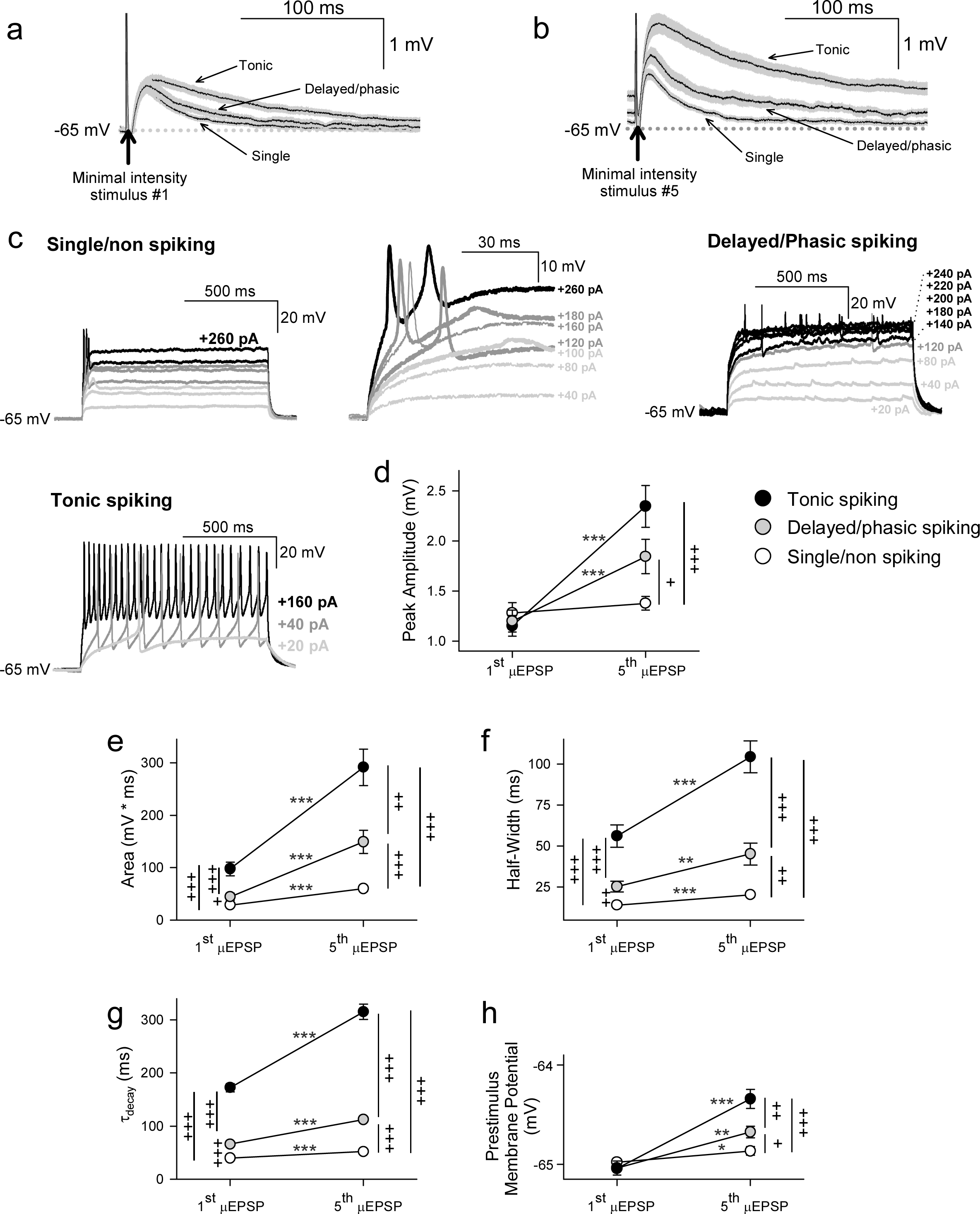
First and fifth μEPSP responses recorded from single/non-spiking, delayed/phasic spiking, and tonic spiking lamina I neuron types. (a) Pure mean (±SEM) μEPSP traces (without synaptic failures) shown for the first (a) and fifth (b) minimal stimulus during the 5 Hz train (arrows indicate stimulus artefacts) for single/non-spiking, delayed/phasic spiking, and tonic spiking lamina I neurons. (c) Representative responses to current injection of each lamina I neuron type including single/non-spiking (inset at right shows magnified view), delayed/phasic spiking, and tonic spiking to current injection (20 pA for 1s). (d) Scatter plots (mean ±SEM) showing μEPSP peak amplitude for the first and fifth mEPSPs from tonic spiking (black symbol), delayed/phasic spiking (gray symbol), and single/non-spiking (white symbol) lamina I neurons (μEPSP #1 vs #5, * p < 0.05, 0** p < 0.01, or *** p < 0.001, paired t-test; + p < 0.05, ++ p < 0.01, or +++ p < 0.001 between lamina I spiking types at μEPSP #1 and #5, t-test), (e) area, (f) half-width, (g) decay, and (h) prestimulus membrane potential.

These results show that primary afferent-lamina I neuron synapses possess distinct kinetic properties that enable μEPSP temporal summation capability (Fig. 4d-h). Importantly, μEPSP summation capacity at a given primary afferent synapse shows correlation with lamina I neuron type (single/non-spiking < delayed/phasic spiking < tonic spiking).

In another approach to elucidate potential μEPSP diversity we applied unsupervised HC analysis to the μEPSP burst dataset. μEPSP kinetic properties including pure first and fifth μEPSP half-width, area, and decay were used for HC which identified three distinct clusters (Fig. 5a-c) referred to as ‘fast,’ ‘intermediate’ and ‘slow’ kinetic properties which we compared to μEPSP classification based on lamina I neuron type (Figs. 5d). Clusters representing first and fifth μEPSP responses from tonic spiking neurons (Fig. 5d) corresponded entirely with clusters representing ‘slow’ μEPSP kinetics for both the first and fifth μEPSP responses (Fig. 5c-e, black symbols) indicative of a homogenous distribution of primary afferent synapses with ‘slow’ μEPSP kinetic properties occurring on tonic spiking lamina I neurons (Fig. 5e). Clusters representing delayed/phasic spiking neurons (Fig. 5d, gray symbols) were affiliated predominantly with ‘intermediate’ clusters for the first and fifth μEPSPs (Fig. 5c-e) while a small portion of μEPSP responses were affiliated with either ‘fast’ (Fig.5c,e) or ‘slow’ (Fig. 5c,e) clusters. Clusters representing single/non-spiking neurons for the first and fifth μEPSPs (Fig. 5d, white symbols) overlapped with both ‘fast’ and ‘intermediate’ kinetics (Fig. 5c-e, white and gray symbols) but not with ‘slow’ kinetics (Fig. 5c-e, black symbols). We conclude that first and fifth μEPSPs can be classified into three major groups consisting of ‘fast’, ‘intermediate’, or ‘slow’ kinetics according to half-width, area, and decay (Fig. 5a-c). Notably, single/non-spiking lamina I neurons contain primary afferent synapses with predominantly ‘fast’ and ‘intermediate’ postsynaptic kinetic properties, while delayed/phasic spiking neurons contain primary afferent synapses with predominantly ‘intermediate’ postsynaptic properties and some with ‘fast’ kinetic properties. Tonic spiking neurons contain primary afferent synapses with exclusively ‘slow’ postsynaptic kinetic properties.

**Figure 5.**
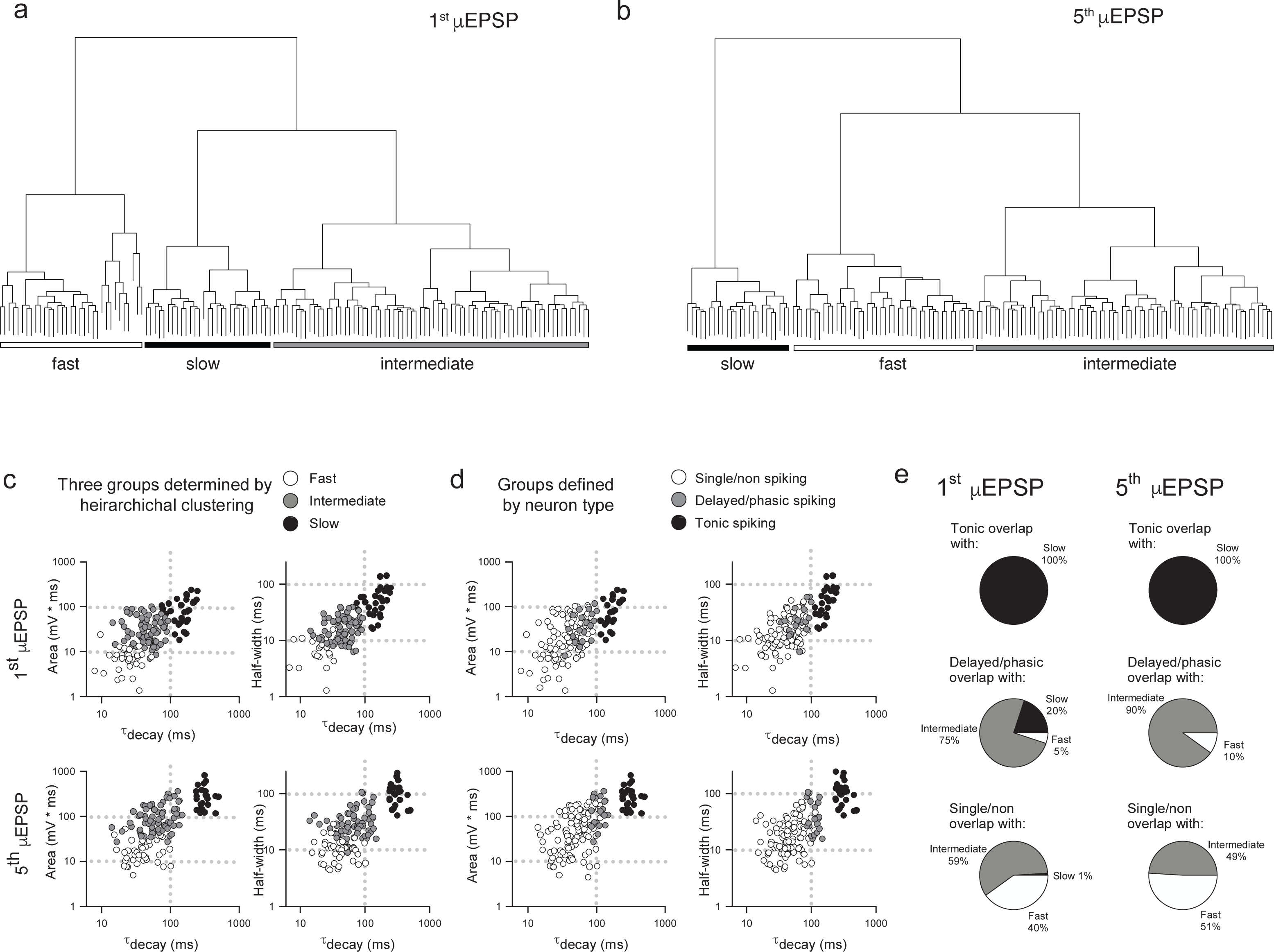
Comparison of μEPSP kinetics by unsupervised hierarchical clustering (HC) and lamina I neuron type. (a) Unsupervised HC analysis of the first (a) and fifth (b) μEPSPs based on half-width, area, and decay identifies three clusters for the first and fifth μEPSPs. (c) Scatter plot of the kinetic data color-coded based on HC-defined subtype affiliation. (d) Scatter plot of the kinetic data color-coded based on lamina I neuron type-defined subtype affiliation. (e) Pie charts summarizing extent of concordance between lamina I neuron type-assigned synapses in panel d and kinetic data-assigned synapses by HC in panel c.

### Link between lamina I neuron type and **μ**EPSP summation capacity is GluN2 subtype-dependent

Having determined that primary afferent-lamina I neuron synapses contain distinct NMDAR GluN2-containing subtypes (Figs. 2-3; Supplementary Figs 2-9), that temporal summation of μEPSPs at these synapses is attributable to NMDAR function (Supplementary Fig. 14), and that HC identified three distinct μEPSP burst clusters, we reasoned that intrinsic properties of μEPSP burst summation capacity derive from distinct primary afferent-lamina I neuron postsynaptic GluN2 subtype-containing NMDARs. To elucidate the role of GluN2 subtype in μEPSP temporal summation, we therefore probed the pharmacological sensitivity of primary afferent-evoked μEPSP bursts recorded from lamina I neurons with known spiking patterns using the selective inhibitor for GluN2A (TCN-201; 3 μM), GluN2B (Ro25-6981; 1 μM), or GluN2D (DQP-1105; 10 μM) administered to aCSF. Baseline μEPSP burst area (mean±SEM) determined from single/non-spiking (88±16 mV*ms), delayed/phasic spiking (115±20 mV*ms), and tonic spiking (295 mV*ms) neurons (Fig. 6d,h,k, respectively) exhibited a progressive increase across neuron type consistent with neuron type-specific temporal summation in our dataset (Fig. 4e). TCN-201 inhibited μEPSP burst responses from single/non- (Fig. 6a,d; n = 4) and delayed/phasic spiking lamina I neuron (Fig. 6f,h; n = 1) but was without effect on a μEPSP burst recorded from a tonic spiking lamina I neuron (Fig. 6i,k; n = 1). Ro25-6981 was also found to inhibit μEPSP burst responses from single/non-spiking (Fig. 6c,d; n = 1) and delayed/phasic spiking (Fig. 6e,h; n = 4) lamina I neurons. However, in experiments in which we tested involvement of GluN2D-containing NMDARs, DQP-1105 was without effect on μEPSP bursts recorded from single/non-spiking (Fig. 6b,d; n = 1) and delayed/phasic spiking (Fig. 6g-h; n = 3) lamina I neuron types but inhibited the μEPSP burst recorded from a tonic spiking lamina I neuron (Fig. 6j-k; n = 1). Our results identify GluN2A and 2B subtype function at primary afferent synapses on single/non-spiking and delayed/phasic spiking lamina I neurons while the GluN2D NMDAR subtype functions predominantly at primary afferent synapses on tonic spiking lamina I neurons. Taken together, we conclude that GluN2 subtypes occur with diversity across different lamina I neuron types and that μEPSP summation at the level of individual primary afferent-lamina I neuron synapses is dependent on distinct GluN2 subtype(s) composition and function at the synapse.

**Figure 6.**
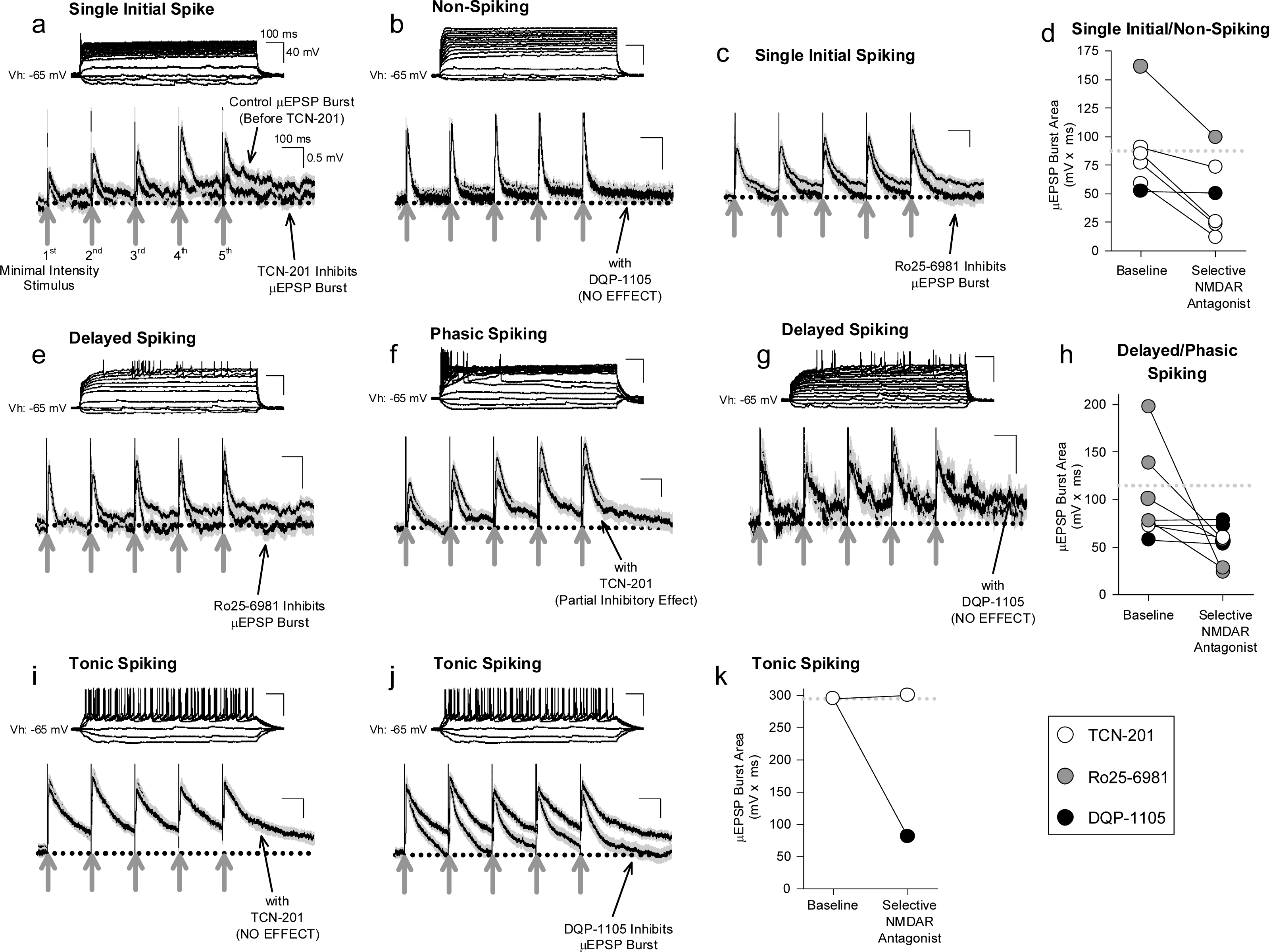
Effect of selective inhibitors for GluN2A (TCN-201), GluN2B (Ro25-6981), or GluN2D (DQP-1105) on representative μEPSP bursts recorded from the three lamina I neuron types. Given the continuous nature of these recordings, these traces include both successes and failures. (a-d) μEPSP bursts recorded from single/non-spiking lamina I neurons exhibited sensitivity to TCN-201 (a) and Ro25-6981 (c), but not to DQP-1105 (b). (d) Summary scatter plot showing μEPSP burst area at baseline and during administration of the selective GluN2 inhibitor to aCSF. (e-h) μEPSP bursts recorded from delayed/phasic spiking lamina I neurons exhibited sensitivity to Ro25-6981 (e) and TCN-201 (f), but not to DQP-1105 (g). (h) Summary scatter plot showing μEPSP burst area at baseline and during administration of the selective GluN2 inhibitor to aCSF. (i-k) μEPSP bursts recorded from a tonic spiking lamina I neuron did not exhibit sensitivity to TCN-201 (i) but exhibited sensitivity to DQP-1105 (j). (k) Summary scatter plot showing μEPSP burst area at baseline and during administration of the selective GluN2 inhibitor to aCSF.

## DISCUSSION

Our strategy to examine single synapse fidelity postsynaptic responses to evoked primary afferent input in the adult rat spinal lamina I subfield identifies this sensory synapse as a major locus of diversity that arises from a heterogenous postsynaptic compartmentalization of single or multiple distinct GluN2 subtypes. We show further that the distinct postsynaptic GluN2 subtype(s) configuration at a given individual afferent synapse on a lamina I neuron defines precisely the computational capability and strength of that synapse.

### Single or multiple GluN2 subtype configurations at primary afferent-lamina I neuron synapses

Principal component and hierarchical analysis combined with selective GluN2 pharmacological testing of μEPSCs recorded from hundreds of individual primary afferent-lamina I neuron synapses identified distinct postsynaptic GluN2 subtype configurations. To our knowledge this is the first demonstration of higher-order assembly of distinct GluN2 subtypes of the NMDAR occurring either individually or in defined combinations at a fully developed, mammalian, glutamatergic synapse. Single-molecule super-resolution imaging in cultured hippocampal neurons has described precise control of the spatial organization of distinct GluN2 subtypes (GluN2A and 2B to date) in defined postsynaptic nanodomains by specific PSD protein constituents ^27^. Here we used an adult, mammalian spinal cord slice preparation to investigate glutamatergic synapses *ex vivo*. Primary afferent-lamina I neuron synapses showing distinctive GluN2-containing postsynaptic localization consisting of either GluN2A alone, GluN2A and 2B, or GluN2B and 2D are indicative of an extensive GluN2 subtype-based heterogeneity of postsynaptic NMDAR organization at a principal sensory glutamatergic synapse.

Previous investigations documenting GluN2 subtype content in spinal lamina I and II have used high intensity stimulation-evoked EPSCs ^44–46^ or spontaneously occurring mEPSCs ^34^ from populations of synapses to assess NMDAR responses. Although these studies provide insight into GluN2 subtype content in the superficial dorsal horn, stimulus protocols that elicit activation of a synapse population are not amenable to elucidating single or multiple GluN2 subtype configurations and function at an individual primary afferent-lamina I neuron synapse. Furthermore, recording miniature or spontaneous unitary synaptic events do not resolve responses in a population of synaptic inputs from responses of an individual synapse. The single-synapse fidelity of our recordings here enabled us to acquire and study defined unitary synaptic events at individual primary afferent-lamina I neuron monosynaptic connections and investigate at high resolution GluN2 subtype diversity and function.

### Functional implications of GluN2 subtype on synaptic computation and transmission in spinal lamina I

Voltage-dependent inward currents mediated by sodium and calcium channels have been proposed to mediate EPSP pattern and integration in the brain ^47^ and spinal cord dorsal horn ^48^. However, the differences in Mg^2+^ block of the distinct μEPSC NMDAR components found here at individual primary afferent-lamina I neuron glutamatergic synapses (Supplementary Fig. 9) not only substantiate the postsynaptic organization of GluN2A, GluN2A and 2B, or GluN2B and 2D we propose to occur at these synapses but highlight their distinct biophysical properties with implications in current flow during membrane depolarization ^28, 39^. We show here that evoked unitary depolarizations at primary afferent-lamina I monosynaptic connections are dependent on both AMPAR- and NMDARs. More specifically, our findings indicate that μEPSP decay and summation throughout the μEPSP burst is shaped primarily by NMDAR-mediated current (Fig. 6, Supplementary Fig. 14) opening the possibility that NMDARs also contribute to the integrative kinetics underlying subthreshold synaptic membrane depolarization ^49^.

In addition to acquiring evoked μEPSP bursts, current injection in our recordings elucidated spiking characteristics of lamina I neurons including single/non-spiking, delayed/phasic spiking, or tonic spiking ^41–43^. This enabled us to classify μEPSP burst properties by neuron spiking type (Fig. 4). However, a definitive correlation could not be made with the three μEPSP clusters categorized by lamina I neuron spiking type and the three clusters of μEPSP kinetics determined by HC analysis (Fig. 5c-e). Though μEPSPs clustered by the tonic spiking lamina I neuron type corresponded entirely with the slower and more prolonged μEPSPs classified by HC analysis, this was not the case for the other two neuron types. Delayed/phasic spiking lamina I neurons exhibited μEPSP kinetic properties that corresponded predominantly with those of intermediate μEPSP kinetics classified by HC while a small percentage of μEPSPs exhibited kinetic properties consistent with fast and slow kinetics defined by HC. Furthermore, μEPSPs evoked at primary afferent-single/non spiking lamina I neurons exhibited kinetic properties that corresponded almost equally with fast and intermediate μEPSP kinetic properties determined by HC. The overall concept that emerges from these results is that μEPSP classification is not exclusively neuron specific but rather synapse specific in that μEPSP kinetics in a given HC-defined group can be found in different lamina I neuron types. μEPSP kinetic properties that are either identical for a given neuron type, ie. neuron-specific, or distinctly different within a given neuron type, ie. synapse-specific, exemplify diversity in the individual μEPSP itself as well as temporal summation capability at primary afferent-lamina I neuron synapses that has major functional implications in subthreshold synaptic computation and strength for lamina I neurons to temporally-patterned sensory input. No other study that we are aware of has demonstrated this degree of heterogeneity in primary afferent-evoked μEPSP responses at individual synapses within and between distinct lamina I neuron types.

The three μEPSP clusters we report here are unexpected because they do not correspond to unitary synaptic depolarization kinetics reported previously in the spinal dorsal horn ^48^. Prescott et al. (2005) found that subthreshold depolarization measured from mEPSPs was similar to that from both delayed/phasic and single spiking lamina I neurons while mEPSPs from tonically spiking lamina I neurons exhibited markedly prolonged depolarization kinetics ^48^, consistent with two classifications of unitary mEPSP kinetics. We do not have a clear explanation for this apparent discrepancy, but the possibility can’t be excluded that the miniature events in the study by Prescott et al. (2005) may have derived from synaptic inputs other than primary afferents. We therefore consider the prospect that mechanisms underlying the kinetics of unitary depolarizations at the primary afferent-lamina I synaptic connection may be distinctly different from those at non-primary afferent synaptic connections. In addition, the number of unitary synaptic recordings obtained in the present study may also merit consideration in that they are relatively high and may therefore favor increased opportunity to detect and decipher previously unrecognized synaptic distinctiveness of NMDAR-dependent biophysical properties of evoked μEPSPs at the primary afferent-lamina I neuron synapse.

The three μEPSC and three μEPSP clusters identified by HC analysis (Fig. 2,5, respectively) support the idea that three distinct GluN2-dependent mechanisms may be at play in mediating different stimulus-evoked membrane depolarization at individual primary afferent-lamina I neuron synapses. The selective GluN2D inhibitor, DQP-1105, was found to be without effect on fast and intermediate μEPSP decay and burst responses in recordings from single/non-spiking and delayed/phasic lamina I neurons (Fig. 6b,d,g-h), but depressed the μEPSP burst exhibiting slow decay kinetics and robust temporal summation recorded from a tonic spiking neuron (Fig. 6j-k). Thus, GluN2D subtype-dependent slow kinetic properties therefore function to broaden μEPSP decay and integrative capacity, and ultimately facilitate burst summation capability to temporally-patterned afferent synaptic input to tonic spiking lamina I neurons. The selective GluN2A inhibitor, TCN-201, was without effect on slow μEPSP decay and summation in a recording from a tonic spiking lamina I neuron (Fig. 6i,k), but depressed μEPSP bursts exhibiting fast decay kinetics and low temporal summation in recordings from single/non- and delayed/phasic spiking neurons (Fig. 6a,d,f,h). GluN2A subtype-dependent fast kinetic properties therefore function to accelerate μEPSP decay and limit integrative capacity to temporally-patterned afferent synaptic input. The selective GluN2B inhibitor, Ro25-6981, depressed μEPSP bursts displaying intermediate kinetic properties in recordings from single/non- and delayed/phasic spiking lamina I neurons (Fig. 6c-e,h) indicative of GluN2B subtype-dependent function in mediating intermediate μEPSP decay and temporal summation. The main concept that emerges from our data is that decay and summation capacity of μEPSPs evoked at a given primary afferent-lamina I neuron synapse are functionally dependent and shaped by the distinct GluN2 subtype synaptic predominance at that synapse which signals to narrowly, intermediately, or widely tune, respectively, the duration of evoked unitary depolarization from resting membrane potential. Ultimately, this would have major implications in subthreshold integrative capacity of different lamina I neuron types to temporally-patterned primary afferent glutamatergic input.

The effects of the selective GluN2 subtype inhibitors examined in our μEPSC recordings revealed that the GluN2 subtype with the longest decay at a given synapse is the main identifiable GluN2 subtype responsible for carrying the greatest charge transfer at that synapse (Supplementary Fig. 6-7). Implicit to these results is that a GluN2 subtype(s) exhibiting residual kinetics identify multiple possible GluN2 subtype combinations localized at a synapse. Taking into account the GluN2A, 2A and 2B, and 2B and 2D synaptic configurations determined in our voltage clamp experiments, we therefore propose the scenario in which primary afferent synaptic connections to single/non-spiking lamina I neurons may contain GluN2A only or GluN2A and 2B postsynaptically, primary afferent synaptic connections to delayed/phasic neurons may contain GluN2A and 2B and some GluN2A only postsynaptically, while primary afferent synapses on tonic spiking lamina I neurons may be comprised exclusively of GluN2B and 2D postsynaptically. Although our data together provide evidence to suggest postsynaptic localization of distinct single or multiple GluN2 subtypes at a given synapse, it remains unclear the functional importance of single or multiple GluN2 subtypes exhibiting differing kinetic properties at a given individual synapse. For instance, in Figure 6f,h, we are limited in our interpretation of whether the depressive effect of TCN-201 on the μEPSP burst is due to inhibition of GluN2-containing NMDARs localized at a given synapse or GluN2-containing NMDARs localized along with GluN2-containing NMDARs at a synapse. We reason that synapse-specific diversity of single or multiple GluN2 subtype combinations postsynaptically endow primary afferent-to-lamina I neuron synapses with single or multiple GluN2 subtype-dependent capability in signaling, modulating membrane potential, and transmission strength.

Single cell transcriptional analysis of neurons in the rodent spinal dorsal horn ^7^ has identified 15 inhibitory (GABAergic) and 15 excitatory (glutamatergic) molecular subtypes of neuron in which Grin2b occurs abundantly in both excitatory and inhibitory subtypes, while Grin2d is low in excitatory and moderately expressed in inhibitory neurons, and Grin2a is found to be low in inhibitory and low to moderate in excitatory neurons. We did not explicitly attempt here to resolve GluN2 subtype function in inhibitory vs excitatory lamina I neurons. We did, however, acquire the spiking pattern of the lamina I neurons recorded in current clamp, as described above. Notably, tonic spiking spinal dorsal horn neurons are generally classified as inhibitory ^41, 42^, but accumulating evidence shows a proportion of this neuron spiking type to be excitatory ^41, 43^. Furthermore, though delayed/phasic spiking dorsal horn neurons have been shown to be inhibitory ^42, 43^, they have also been shown to be predominantly excitatory ^41–43^.

Similarly, though a small proportion of single/non-spiking dorsal horn neurons have been found to be inhibitory ^42, 43^, this neuron type has also been found to be predominantly excitatory ^41–43^. Notwithstanding these apparent discrepancies, the synapse-specificity we propose here rooted in distinct GluN2 subtype configurations and computation capability may represent a basis on which multiple CNS mechanisms (eg. excitatory and inhibitory) are driven by glutamatergic synaptic input and tightly modulated by distinct postsynaptic GluN2 subtype configurations.

### Synaptic GluN2 subtype heterogeneity as a determinant of transmission fidelity in the CNS

We used the *ex vivo* primary afferent-lamina I neuron synaptic preparation as a model glutamatergic circuit in which single-synapse fidelity postsynaptic responses were investigated. Based on our results showing synapse-specific GluN2 subtype heterogeneity across individual glutamatergic synapses, we propose for consideration major roles of GluN2 subtype diversity in neuron excitability throughout the CNS. The functional dependence of μEPSP shape and μEPSP burst summation capacity to temporally patterned input on the diverse single or multiple GluN2 subtype configurations with differing kinetics at a given synapse would be predicted, in an ensemble of glutamatergic inputs, to impact EPSP-spike coupling ^50^ by gauging membrane potential depolarization and, ultimately, the neuron reaching spiking threshold. We also put forward here the concept that GluN2 protein subtype heterogeneity in synaptic ensembles to a given neuron may serve a coding function that coordinates the synaptic integrative fidelity of that neuron to repetitive input activity (coincidence detection) which, importantly, would further influence neuron firing.

The NMDAR dendritic spike is the dominant mechanism by which dendritic integration of glutamatergic inputs through the recruitment of NMDAR channels leads to firing of a neuron ^51, 52^. Since its discovery in cortical pyramidal cell dendrites over 20 years ago ^53^ the NMDAR spike continues to receive considerable focus in different cortical regions ^54–58^. Notably, recent evidence has uncovered critical features of NMDAR-containing synapses that contribute to the generation of spikes including synapse number ^59^, proximity to other synapses ^59^, spatial clustering ^60–62^, as well as their dendritic location ^63–65^. Given our findings here, we propose for consideration the potential role GluN2 synaptic diversity and organization may impose on dendritic spike threshold and efficacy in firing in neurons in which NMDAR spikes predominate.

Key distinctions in structural features of individual glutamatergic synapses including spine size, volume ^66–70^, and PSD area ^71–75^ show correlation with physiological transmission strength. Whether or not proteomic complexity at synapses including that of GluN2 synaptic heterogeneity may be causative, correlative, or independent of PSD or synapse size, we suggest that synaptic distinctiveness, rooted in GluN2 subtype diversity, is a critical determinant in grading subthreshold glutamatergic synaptic strength.

Molecular and architectural complexity across glutamatergic synapses, arising from a multitude of differences in postsynaptic proteome constituents and their nanostructure organization ^3, 5, 6^, are beginning to emerge as determinants of individual synaptic diversity in the brain ^1^ and spinal dorsal horn ^8^. We demonstrate here that a major locus of proteome complexity exists in the spinal lamina I nociceptive subfield and involves extensive postsynaptic GluN2 subtype diversity at the primary afferent-lamina I neuron synapse. Collectively, our data are consistent with a model wherein synapse-specific organization of distinct GluN2 subtypes serves as a fundamental subthreshold coding mechanism and a core determinant of how physiological sensory information is processed and transferred from the periphery to the brain. Future studies that scrutinize NMDAR subtype diversity and function at this principal sensory synapse in models of neuropathy and inflammation will undoubtedly advance our understanding of synaptic mechanisms underlying chronic pain associated with trauma or disease. Our findings also offer insight into complexity of glutamatergic synapses and their function in other regions of the CNS in health and disease.

Supplementary Figure 1. (a) Scatter plot illustrating minimal stimulation-evoked μEPSCs (Vh: -65 mV) eliminated by TTX (500 nM). (b) Left: Sample traces showing individual traces (Vh: -65 mV), (*i-v*) from (a), demonstrating successful μEPSCs and failures in response to minimal stimulation (the stimulus artefact is depicted by the gray arrow). A spontaneous EPSC is shown in trace (*iii*). Right: Successful μEPSCs are blocked following TTX administration. mEPSCs are shown in traces (*vi-ix*). (c) Scatter plot showing evoked μEPSCs (Vh: +60 mV) at minimal stimulation threshold (gray arrow at left; sample μEPSCs shown at *i-ii*), and slightly above threshold (thick gray arrow at right) which evoked a second level of all-or-none responses (sample μEPSCs shown at *iii-iv*). Stimulus intensity below that of minimal stimulation intensity (thin gray arrow at time point 3 min) did not evoke μEPSCs. (d) Paired-pulse minimal stimulation experiment showing average amplitude (± SEM) of μEPSCs (Vh: +60 mV) to the second stimulus (d(*ii*)) was greater than that of the first (d(*i*)) when the first stimulus also evoked a μEPSC. The average amplitude of the μEPSCs to the second stimulus (d(*iii*)) that followed a failure of response to the first stimulus was also identical, if not greater than, the average amplitude (± SEM) of the response to the second stimulus that followed the first stimulus non-failures. Right: d(*i*) scatter plot showing μEPSC amplitude with a single distribution (insert below) evoked during the first stimulus. d(*ii*) and d(*iii*) show single distributions of μEPSCs to the second stimulus. (e) Same experiment as in (d) but recorded from a different lamina I neuron. Insert above shows that single pulse minimal stimulation evoked a mean (± SEM) μEPSC recorded prior to paired-pulse stimulation (left) is identical to the mean (± SEM) μEPSC recorded in the same experiment but after paired-pulse stimulation (right).

Supplementary Figure 2. (a) Bi-plot representing the individual data points of the full experimental dataset in the PCA space of the variables in the study. The first two dimensions describe 60% of the dataset variance. (b) Heatmap representing the contributions of variables in accounting for the variability in the PCA components. Values are expressed in percentage of the total variability in the PCA components. (c) Scatter plot of the raw data color-coded based on HC-defined subtype affiliation.

Supplementary Figure 3. Representative fast, intermediate, and slow μEPSCs. (a) Sample trace (black) showing an individual fast decay μEPSC (Vh: +60 mV) from a representative spinal lamina I neuron (gray arrow indicates stimulus artefact). The gray trace shows a synaptic failure. (b) Scatter plot of successful synaptic μEPSCs responses (Vh: +60 mV) and failures over time (thick gray bar) evoked by minimal stimulation (0.2 Hz). The thin gray arrow indicates subthreshold stimulation intensity while the thick gray arrow indicates threshold at which μEPSCs can be evoked. (c) NMDAR amplitude histogram showing relative distribution for successful μEPSCs in this recording (a-b), fit with a single Gaussian function. (d) Averaged traces (± SEM) showing successful μEPSCs held at +60 mV in this recording (a-c). (e) Single exponential fitting of the decay component of the averaged μEPSC in (d) held at +60 mV. (f-j) Recording of a representative intermediate decay μEPSC (Vh: +60 mV) described as per (a-e). (k-o) Recording of a representative slow decay μEPSC (Vh: +60 mV) described as per (a-e).

Supplementary Figure 4. Multiple representative traces of fast (a-b), intermediate (c-d), and slow (e-f) μEPSCs (from recordings in Supplementary Figure 3) illustrating stable latency, amplitude, and decay.

Supplementary Figure 5. D-APV blocks fast, intermediate, and slow μEPSC decay components. (a) Averaged traces (± SEM) from a representative recording showing successful fast decay μEPSCs (Vh: +60 mV) before (baseline) and with D-APV (100 mM in aCSF). Averaged traces (± SEM) from the same recording showing successful μEPSCs held at -65 mV. (b) Averaged traces (± SEM) from a representative recording showing successful intermediate decay μEPSCs (Vh: +60 mV) before (baseline) and with D-APV. Averaged traces (± SEM) from the same recording showing successful μEPSCs held at -65 mV. (c) Averaged traces (± SEM) from a representative recording showing successful slow decay μEPSCs (Vh: +60 mV) before (baseline) and with D-APV. Averaged traces (± SEM) from the same recording showing successful μEPSCs held at -65 mV with D-APV present.

Supplementary Figure 6. Effect of DQP-1105 (10 μM mM in aCSF) on the decay and charge transfer components of fast, intermediate, and slow μEPSCs. (a) Averaged traces (± SEM) from a representative recording showing slow decay μEPSCs (Vh: +60 mV) before (baseline) and with DQP-1105 (gray arrow indicates stimulus artefact). (b) Averaged traces (± SEM) from a representative recording showing intermediate decay μEPSCs (Vh: +60 mV) before (baseline) and with DQP-1105. (c) Averaged traces (± SEM) from a representative recording showing fast decay μEPSCs (Vh: +60 mV) before (baseline) and with DQP-1105. (d) Summary scatter plot showing effect of DQP-1105 on μEPSC charge transfer from recordings before (baseline) and with DQP-1105. (e) Summary histogram showing a depressive effect of DQP-1105 on charge transfer of representative slow decay μEPSCs from (d) (n = 3; p < 0.05 vs baseline, paired t-test). (f) Summary histogram showing no effect of DQP-1105 on charge transfer of representative intermediate decay μEPSCs from (d) (n = 9; p > 0.05 vs baseline). (g) Summary histogram showing no effect of DQP-1105 on charge transfer of representative fast decay μEPSCs from (d) (n = 5; p > 0.05 vs baseline).

Supplementary Figure 7. Effect of Ro25-6981 (1 μM in aCSF) on the decay and charge transfer components of fast, intermediate, and slow μEPSCs. (a) Averaged traces (± SEM) from a representative recording showing slow decay μEPSCs (Vh: +60 mV) before (baseline) and with Ro25-6981 (gray arrow indicates stimulus artefact). (b) Averaged traces (± SEM) from a representative recording showing intermediate decay μEPSCs (Vh: +60 mV) before (baseline) and with Ro25-6981. (c) Averaged traces (± SEM) from a representative recording showing fast decay μEPSCs (Vh: +60 mV) before (baseline) and with Ro25-6981. (d) Summary scatter plot showing effect of Ro25-6981 on μEPSC charge transfer from recordings before (baseline) and with Ro25-6981 present. (e) Summary histogram showing a partial depressive effect of Ro25-6981 on charge transfer of representative slow decay μEPSCs from (d) (n = 5; p < 0.001 vs baseline, paired t-test). (f) Summary histogram showing a depressive effect of Ro25-6981 on charge transfer of representative intermediate decay μEPSCs from (d) (n = 4; p < 0.01 vs baseline, paired t-test). (g) Summary histogram showing no effect of Ro25-6981 on charge transfer of representative fast decay μEPSCs from (d) (n = 3; p > 0.05 vs baseline).

Supplementary Figure 8. Effect of TCN-201 (3 μM in aCSF) on the decay and charge transfer components of fast μEPSCs. (a) Averaged traces (± SEM) from a representative recording showing fast decay μEPSCs (Vh: +60 mV) before and with TCN-201 (gray arrow indicates stimulus artefact). (b) Summary scatter plot showing effect of TCN-201 on μEPSC charge transfer (Vh: +60 mV) from recordings before (Baseline) and with TCN-201 present. (c) Summary histogram showing depressive effect of TCN-201 on charge transfer of fast decay μEPSCs from (b) (n = 3; p < 0.01 vs baseline, paired t-test). (d) Mean (±SEM) charge transfer of fast μEPSCs (Vh: +60 mV) from Figure 3a, represented by (*i*), is not significantly different from mean (±SEM) charge transfer of baseline μEPSCs held at +60 mV from panel ‘a’, represented by (*iii*). Mean (±SEM) charge transfer of μEPSCs held at -65 mV from Figure 3a, represented by (*ii*) and considered to be mediated predominantly by AMPAR-mediated charge transfer, is not significantly different from mean (±SEM) charge transfer of baseline μEPSCs held at +60 mV from panel ‘b’ with TCN-201 present in aCSF, represented by (*iv*). (e) We reasoned that the difference in charge transfer between (*i*) and (*ii*) is the pure fast NMDAR component (white bar), and that the difference in charge transfer between (*iii*) and (*iv*) is the pure NMDAR component mediated by GluN2A (red bar). The lack of difference (p > 0.05) is therefore interpreted to indicate that the baseline NMDAR component of the fast μEPSC (panels a-b) is mediated exclusively by GluN2A.

Supplementary Figure 9. Summary of kinetics of μEPSCs recorded at the holding potential of - 65 mV (from Figure 3). (a) Late component of μEPSCs (Vh: -65 mV) exhibited a significant increase in half-width (p < 0.01, unpaired, two-tailed t-test) and charge transfer (p < 0.05, 0.01, 0.001, unpaired, two-tailed t-test) across the three groups. Dotted lines in box plots indicates mean. (b) Percent inhibition of the amplitude of the NMDAR component of μEPSCs held at -65 mV in fast, intermediate, and slow decay groups from Figure 3. The percent inhibition of the NMDAR component of μEPSCs from the ‘fast’ group was similar to that of μEPSCs from the ‘intermediate’ group, whereas the percent inhibition of the NMDAR component of μEPSCs from the ‘slow’ group was less. Percent inhibition of the amplitude of the NMDAR component of μEPSCs held at -65 mV was calculated by dividing the difference between mean amplitude of the NMDAR component of μEPSCs held at -65 mV and the mean amplitude of the NMDAR component of μEPSCs held at +65 mV by the mean amplitude of the NMDAR component of μEPSCs held at +65 mV and multiplying the quotient by 100.

Supplementary Figure 10. Lack of effect of NMDAR inhibitors on μEPSC failure rate. (a) Averaged traces (± SEM) from a representative recording (at left) showing μEPSCs (Vh: +60 mV) before (baseline) and with D-APV (right). Gray arrow indicates stimulus artefact. Below: Scatter plot of successful synaptic μEPSCs responses (Vh: +60 mV) and failures over time evoked by minimal stimulation (0.2 Hz). (b) Scatter plot showing failure rate (mean ±SEM) during baseline and with administration of D-APV, TCN-201, Ro25-6981, or DQP-1105 in experiments in which these inhibitors were tested (n = 46, p > 0.05, paired t-test). (c) Scatter plot showing failure rate (mean ±SEM) during baseline and with administration of D-APV only (n = 11, p > 0.05, paired t-test).

Supplementary Figure 11. μEPSCs and μEPSPs recorded sequentially in the same lamina I neuron during a brief train composed of 5 stimuli delivered every 200 ms (5 Hz) at 5 s intervals (0.2 Hz) at minimal stimulation intensity in voltage clamp and then current clamp modes, respectively. (a) Mean (±SEM) μEPSC (Vh: -65 mV) trace showing continuous current clamp recording during the 5 Hz train (arrows indicate stimulus artefacts throughout the train). Given the continuous nature of these recordings, these traces include both successes and failures. (b) Scatter plots (mean ±SEM; n = 99) showing μEPSC peak amplitude, half-width, charge transfer (p < 0.001, non-parametric repeated measures, μEPSC #1 vs #5, p < 0.001, t-test), and rise time during the 5 Hz train. (c) Mean (±SEM) μEPSP trace showing continuous voltage clamp recording during the 5 Hz train (arrows indicate stimulus artefacts throughout the train). Given the continuous nature of these recordings, these traces include both successes and failures. (d) Scatter plots (mean ±SEM; n = 100) showing μEPSP peak amplitude (non-parametric repeated measures, p < 0.001, t-test; μEPSP #1 vs #5, p < 0.001), half-width (non-parametric repeated measures, p < 0.001, t-test; μEPSP #1 vs #5, p < 0.001), area (non-parametric repeated measures, p < 0.001, t-test; μEPSP #1 vs #5, p < 0.001), and rise time (p > 0.05) during the 5 Hz train.

Supplementary Figure 12. Pure μEPSCs (successes only, data extracted from Supplementary Figure 11a) shown at each stimulus during the 5 Hz train at minimal stimulation intensity in voltage clamp (Vh: -65 mV). (a) Pure mean (±SEM) μEPSC traces (without synaptic failures) shown at each minimal stimulus (arrows indicate stimulus artefacts throughout the train). Representative synaptic failures depicted by the flat traces (mean ±SEM) are shown at each minimal stimulus. (b) Scatter plots (mean ±SEM) showing μEPSC peak amplitude (p > 0.05, μEPSC #1 vs #5), half-width (p > 0.05, μEPSC #1 vs #5), charge transfer (p > 0.05, μEPSC #1 vs #5), rise time (p > 0.05, μEPSC #1 vs #5), and failure rate (p < 0.05, non-parametric repeated measures, μEPSC #1 vs #5, p < 0.05, t-test) for each minimal stimulus.

Supplementary Figure 13. Pure μEPSPs only (successes only, data extracted from Supplementary Figure 11c) shown at each stimulus during the 5 Hz train at minimal stimulation intensity in current clamp. (a) Pure mean (±SEM) μEPSP traces (without synaptic failures) shown at each minimal stimulus during the 5 Hz train (arrows indicate stimulus artefacts throughout the train). (b) Scatter plots (mean ±SEM) showing μEPSP peak amplitude (p < 0.001, repeated measures ANOVA; μEPSP #1 vs #5, p < 0.001, t-test), area (p < 0.001, μEPSP #1 vs #5, p < 0.001, statistics as per above), half-width (p < 0.001, μEPSP #1 vs #5, p < 0.001, statistics as per above), rise time (p > 0.05, μEPSP #1 vs #5), prestimulus membrane potential (p < 0.001, μEPSP #1 vs #5, p < 0.001, statistics as per above), decay (p < 0.001, μEPSP #1 vs #5, p < 0.001, statistics as per above), and failure rate (p < 0.05, μEPSP #1 vs #5, p < 0.05, statistics as per above) for each minimal stimulus.

Supplementary Figure 14. Effects of D-APV or MK-801 on μEPSP bursts. (a) Mean (±SEM) μEPSP burst traces showing continuous recording during the 5 Hz train (gray arrows indicate stimulus artefacts throughout the train) during baseline (summated mean±SEM trace), with D-APV (100 mM), followed by CNQX (10 mM). Given the continuous nature of these recordings, these traces include successes and failures. (b) Mean (±SEM) μEPSP burst traces showing continuous recording during the 5 Hz train (arrows indicate stimulus artefacts throughout the train) during baseline (summated mean±SEM trace), with the NMDAR blocker, MK-801 (10 μM), followed by CNQX (10 mM).

## Methods

### Animals

Experiments were carried out using adult, male Sprague Dawley rats (from Charles River, P70-80, 325-350g), and approved by the Hospital for Sick Children Animal Care Committee and in accordance with policies of the Canadian Council on Animal Care.

### Spinal cord isolation and slice preparation

The lumbar spinal cord was isolated from anesthetized rats (urethane administered by intraperitoneal injection, 20% (wt/vol)) by rapid dissection through ventral laminectomy and placed immediately in ice-cold protective artificial cerebralspinal fluid (paCSF) solution containing (in mM): 82 NaCl, 3 KCl, 1.25 NaH_2_PO_4_, 9.3 MgCl_2_, 24 D(+)-glucose, 20 NaHCO_3_, 0.1 CaCl_2_ and 70 sucrose (chemicals from Sigma Millipore) saturated in carbogen (95% O_2_, balance 5% CO_2_; pH 7.40 and 315-325 mOsm). After removal of the meninges and ventral roots under a dissecting microscope (Leica MS 5), the ventral surface of the lumbar segment was glued (Vetbond, 3M) to an agar block (4% agarose, low melting point, analytical grade, Promega, Fisher Scientific; 4% agarose was prepared using raCSF(see below)) in a vertical orientation which in turn was glued in place in a vibratome (Leica VT1000 S) slicing chamber containing ice-cold, carbogen-saturated paCSF for slice preparation. Following sectioning (blade advance speed of ∼ 0.02-0.03 mm/s and lateral amplitude of ∼2.5 mm), transverse spinal cord slices (500-600 μm) containing the dorsal rootlets were transferred to incubation aCSF (iaCSF) containing (in mM): 90 NaCl, 3 KCl, 1.25 NaH_2_PO_4_, 9.7 MgCl_2_, 24 D(+)-glucose, 19 NaHCO_3_, 0.1 CaCl_2_ and 60 sucrose saturated with carbogen (pH 7.40, 315-325 mOsm) and maintained at 30±1°C until recording.

### Electrophysiology and data acquisition

For data acquisition, a single transverse spinal cord slice was transferred to the recording chamber and superfused (2 ml/min) with recording artificial cerebrospinal fluid (raCSF) composed of (in mM): 126 NaCl, 3 KCl, 1.25 NaH_2_PO_4_, 2 MgCl_2_, 24 D(+)-glucose, 24 NaHCO_3_ and 2 CaCl_2_ saturated with carbogen at 28-30°C (pH = 7.40; 315-325 mOsm).

Whole cell recordings were from neuronal cell bodies (visualized method (Zeiss Axioskop 2FS microscope) located in the darker, striated region (lamina I) ventral to the outer white matter tracts and dorsal to the substantia gelatinosa, the translucent laminar region that demarcates lamina II ^34^ and were carried out using patch pipettes (5-6 MΩ) containing Cs-gluconate intracellular solution (ICS) composed of (in mM): 105 Cs-gluconate, 17.5 CsCl, 10 HEPES, 10 BAPTA (Life Technologies), 5 QX-314 (Millipore Sigma), 2 Mg-ATP, 0.3 GTP (pH = 7.20; 290 mOsm) or K-gluconate ICS composed of (in mM): 132.5 K-gluconate, 17.5 KCl, 10 HEPES, 0.2 EGTA, 2.0 Mg-ATP, 0.3 GTP (pH 7.20, 290 mOsm). Patch pipettes were pulled from filament-containing borosilicate glass capillaries (TW150F-4; World Precision Instruments Inc., Sarasota, FL, USA) using a Flaming/Brown Micropipette Puller (Model P-97, Sutter Instruments Co., USA), the tips fire-polished using a micro forge (model Micro Forge MF-830, Nareshige; Tokyo, Japan), and positioned for recording using a MP-225 Sutter micromanipulator (Sutter Instrument Company).

Data acquired from lamina I neurons included unitary excitatory postsynaptic excitatory postsynaptic currents (μEPSCs) or unitary excitatory postsynaptic potentials (μEPSPs) evoked by stimulating a single primary afferent synaptic connection to the lamina I neuron recorded. To isolate activation of an individual synaptic connection between a primary afferent axon and a lamina I neuron, the tip of an in-house designed and constructed (by GMP) fine bipolar stimulating electrode (consisting of two ∼50 μm diameter single nichrome wire electrodes (A-M Systems, Inc.) separated by ∼15 μm formvar insulation coating) was placed in the dorsal rootlet entry zone which contains axons of primary afferent neurons that are monosynaptically connected to lamina I neurons. Activation of a putative individual axon at the electrode tip consisted of single pulses of electrical stimulation (0.08 ms duration at 0.2 Hz delivered to the electrode via an S48 Grass Stimulator) with stimulus intensity increased gradually beginning at 0 stimulus intensity (minimal stimulation) until all-or-none unitary postsynaptic responses of the lamina I neuron recorded were detected at which point stimulus intensity (i.e. stimulus threshold) was no longer adjusted (see Results).

μEPSC recordings acquired from lamina I neurons were voltage-clamped at +60 mV using pipettes containing Cs-gluconate ICS. In most of these voltage-clamp recordings μEPSCs were also acquired at a holding potential of -65 mV. μEPSPs were also acquired from lamina I neurons resting at -65 mV using pipettes containing K-gluconate ICS. Of note, prior to acquiring μEPSPs in a given recording we acquired first in the same recording μEPSCs in voltage-clamp mode (Vh: -65 mV). Rationale and further details of these experiments are provided in the Results.

In all experiments, raCSF contained [R-(*R*\*,*S*\*)]-5-(6,8-Dihydro-8-oxofuro[3,4-*e*]-1,3-benzodioxol-6-yl)-5,6,7,8-tetrahydro-6,6-dimethyl-1,3-dioxolo[4,5-*g*]isoquinolinium chloride (bicuculline methochloride, 10 μM dissolved in H_2_O; Tocris Bioscience). The following compounds were added to raCSF as indicated in the Results section: D-(-)-2-Amino-5-phosphonopentanoic acid (D-APV, 100 μM dissolved in H2O; Tocris Bioscience), (5*S*,10*R*)-(+)-5-Methyl-10,11-dihydro-5*H*-dibenzo[*a*,*d*] cyclohepten-5,10-imine maleate (MK-801, 10 μM dissolved in H_2_O; Tocris Bioscience), 3-Chloro-4-fluoro-*N*-[4-[[2-(phenylcarbonyl) hydrazino]carbonyl]benzyl]benzene-sulfonamide (TCN 201, 3 μM dissolved in DMSO; Tocris Bioscience), (α*R*,β*S*)-α-(4-Hydroxyphenyl)-β-methyl-4-(phenylmethyl)-1-piperidinepropanol maleate (Ro25-6981, 1 μM dissolved in H_2_O; Tocris Bioscience), 5-(4-Bromophenyl)-3-(1,2-dihydro-6-methyl-2-oxo-4-phenyl-3-quinolinyl)-4,5-dihydro-γ-oxo-1*H*-pyrazole-1-butanoic acid (DQP 1105, 10 μM dissolved in DMSO; Tocris Bioscience), or 6-Cyano-7-nitroquinoxaline-2,3-dione disodium (CNQX, 10 μM dissolved in H_2_O; Tocris Bioscience), or tetrodotoxin (TTX, 0.5 μM dissolved in H_2_O; Alomone Labs). Recordings were carried out with the experimenter (GMP) blind to the NMDAR GluN2-selective and nonselective NMDAR inhibitors/blockers tested.

Raw data were amplified using a MultiClamp 700A amplifier and a Digidata 1322A acquisition system sampled at 10 kHz and low-pass filtered at 2.4 kHz, and analyzed with Clampfit 10.3 (Axon Instruments), Microsoft Office Excel, and Sigmaplot 11 software (Systat Software, Inc.). Junction potential was accounted for in all current-clamp experiments. Data from voltage-clamp experiments were discarded if the series resistance monitored in the recording changed by more than 15%.

### Data analysis

Data from minimal stimulation experiments were considered *bona fide* minimal stimulation-elicited unitary postsynaptic responses and used for analysis only if the unitary responses in a given experiment exhibited a clear stimulus threshold (all-or-none postsynaptic events) determined upon increasing gradually stimulation intensity from 0 to the threshold stimulus intensity, a clear unimodal, normal distribution of the events, as well as a consistent latency with each stimulus (see Results). All recordings in every experiment were inspected visually to verify failure and non-failure responses. μEPSC kinetic parameters measured for analysis (including decay, amplitude, half-width, charge transfer, and rise time) were calculated from the mean μEPSC trace of all the non-failure responses in that experiment. The decay time constant(s) of the mean μEPSC (Vh: +60 mV) was calculated in Clampfit by least-squares fitting with one or more exponentials. The best-fit of a given μEPSC decay was determined by comparing *R*^2^ for the respective fit(s) of that decay. In current-clamp experiments, μEPSP kinetic parameters including peak amplitude, half-width, and area under the curve were also measured (see Results for details). Failure rate (%) of unitary events in a given minimal stimulation recording was calculated as the percentage of failures that occurred in that given recording.

### Statisitics

Comparisons between independent groups were carried out by two-tail t-test (no equal variance assumed). Comparisons between paired groups were carried out by paired t-test, comparisons on repeated measures were performed by repeated measures ANOVA. Significance was established at α=0.05, or at a corrected value following Bonferroni correction for multiple comparisons where appropriate. Kinetic data were normalized, scaled and analyzed in R by PCA and K-means clustering, followed by gap-statistic to select the optimal number of clusters. HC was built using the euclidean distance method and the ward.d2 linkage method.

## Acknowledgements

This study was supported by a grant from Canadian Institutes of Health Research (CIHR, FDN-154336) and by the Krembil Foundation to MWS. We thank L.Y. Wang (The Hospital for Sick Children) for critical reading of the manuscript.

## Notes

### Competing Interest Statement

The authors have declared no competing interest.

