## Supplementary Figures for "Synapse-specific diversity of distinct postsynaptic GluN2 subtypes defines transmission strength in spinal lamina I"

### Pitcher et al., Supplementary Figure 1

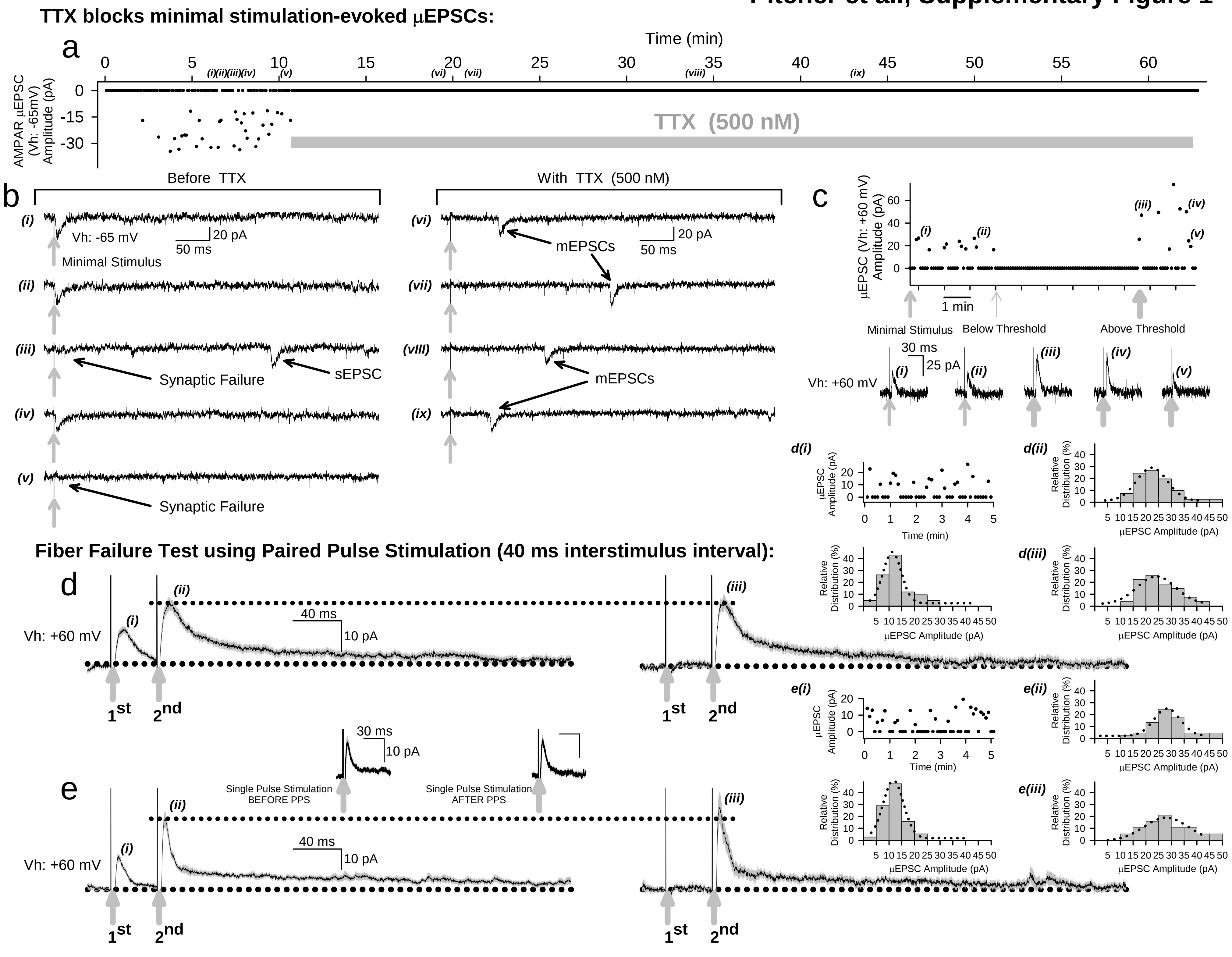

Pitcher et al., . Supplementary Figure 2

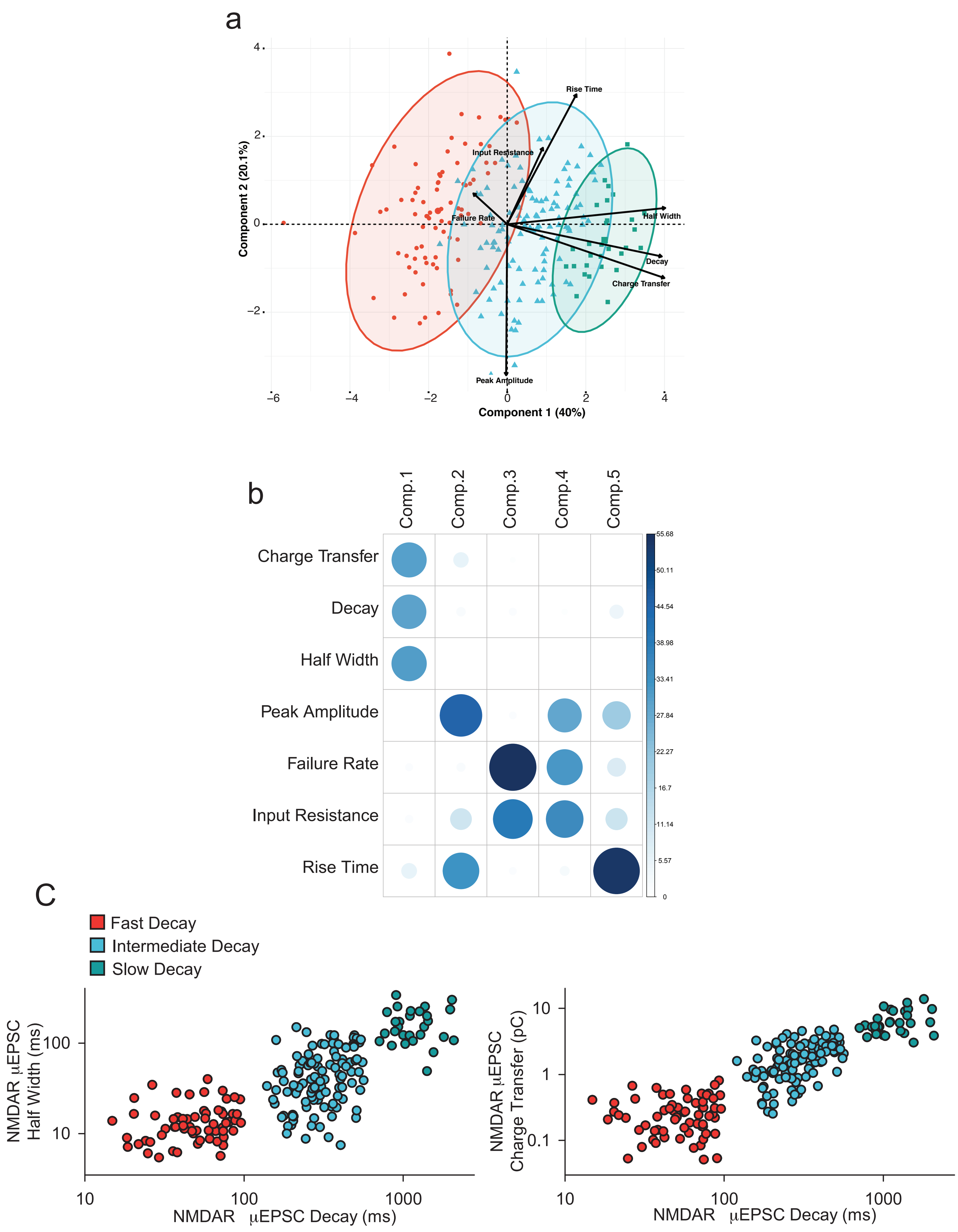

### Pitcher et al., Supplementary Figure 3

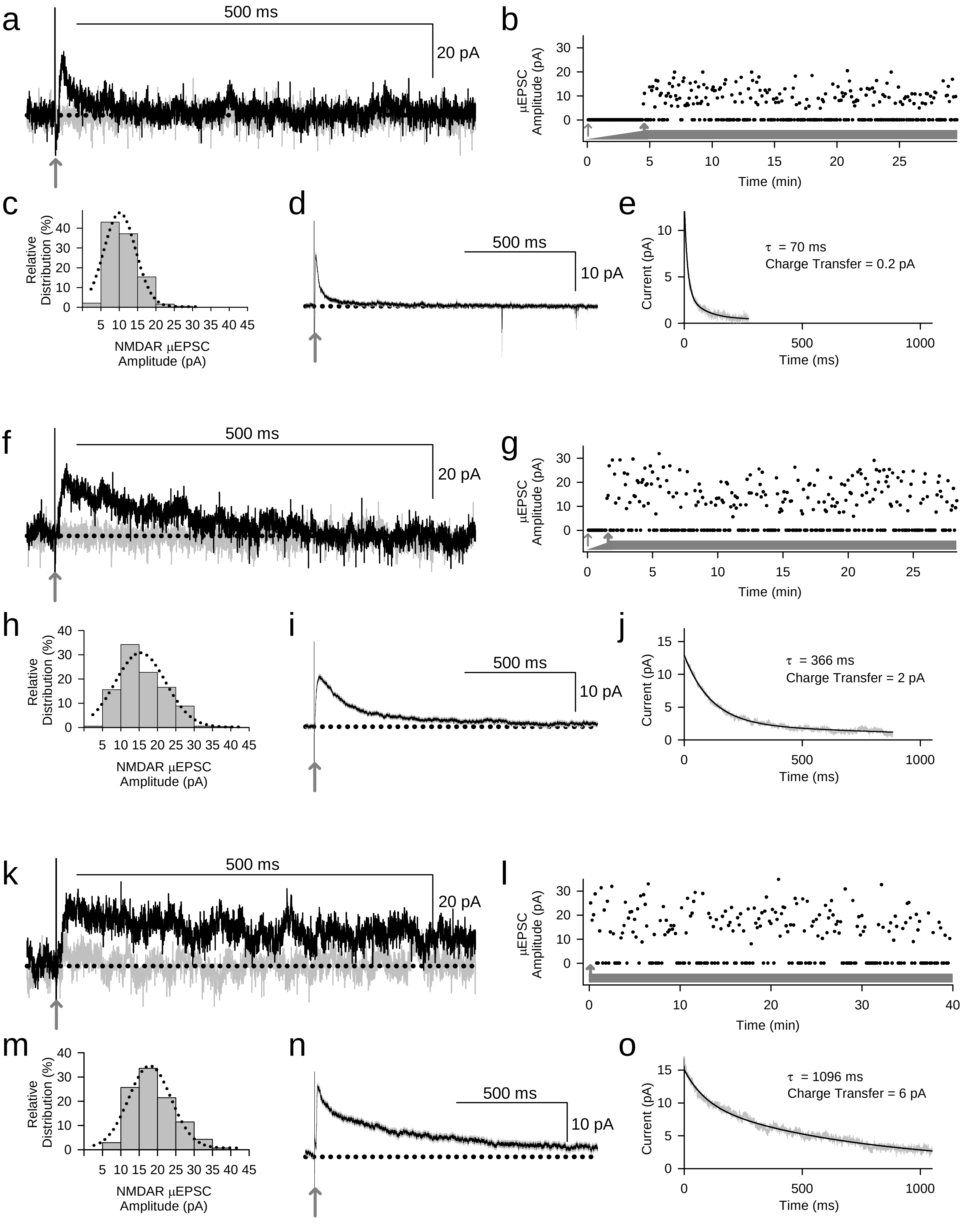

**Pitcher et al., Supplementary Figure 4**

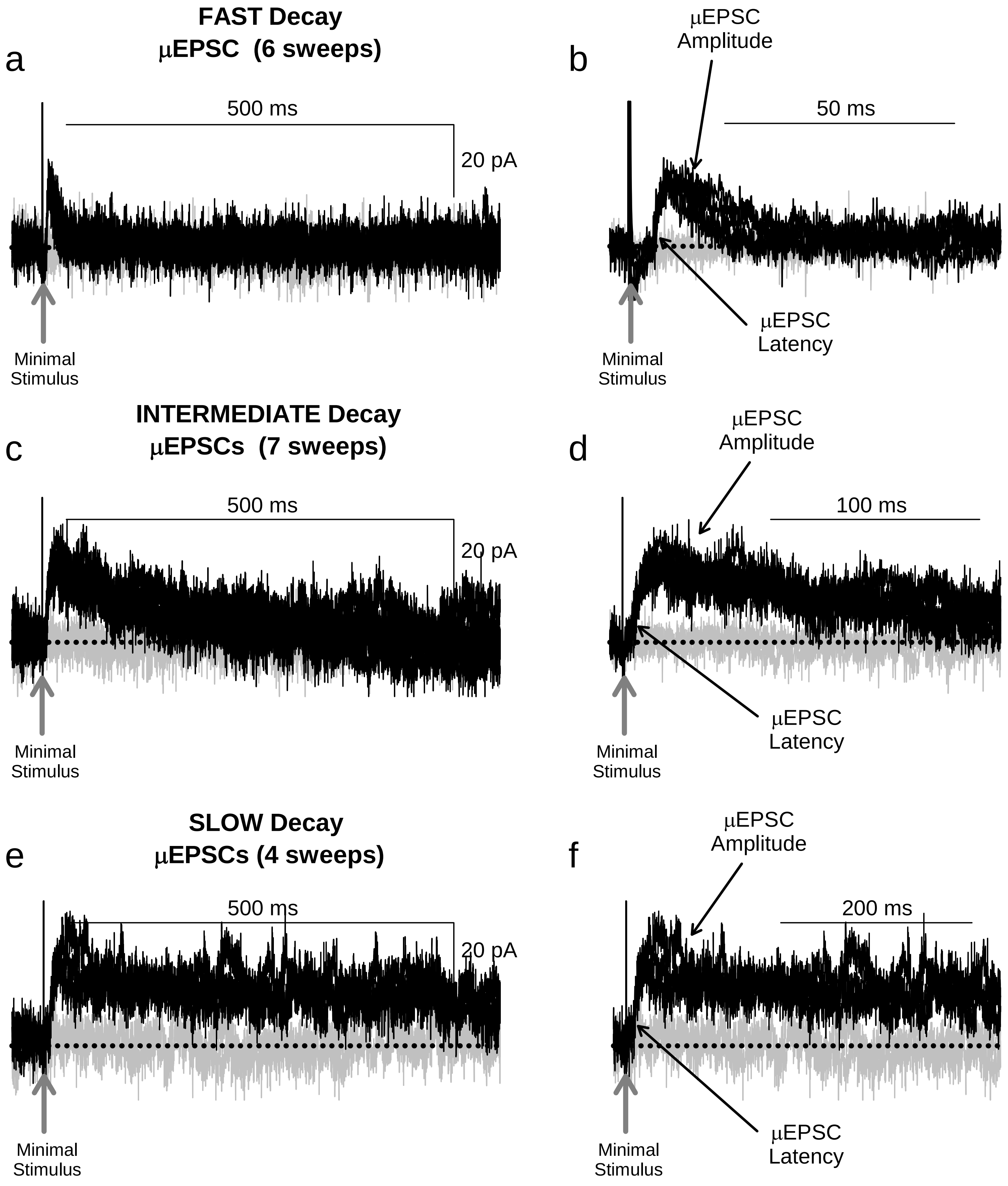

### Pitcher et al., Supplementary Figure 5

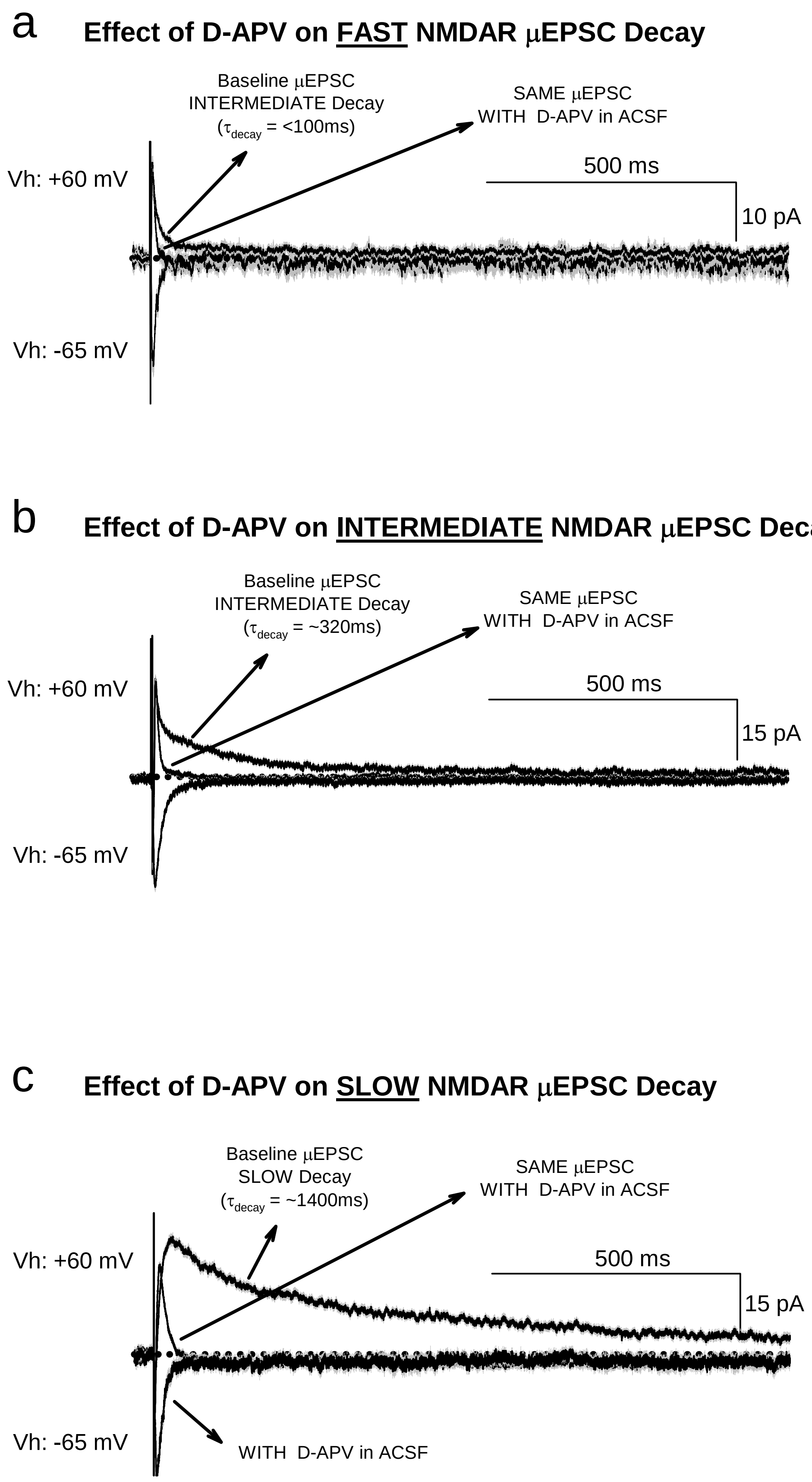

### Pitcher et al., Supplementary Figure 6

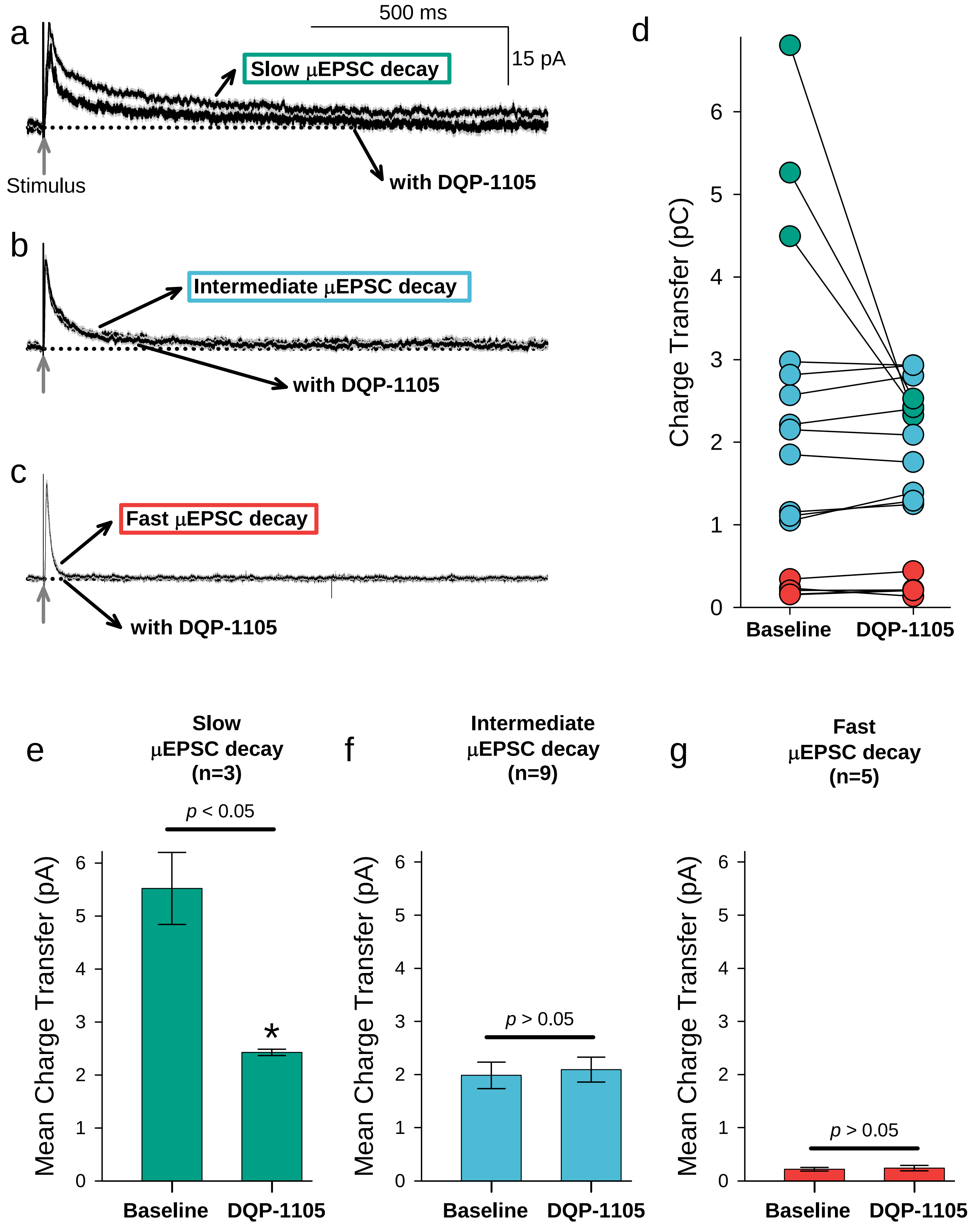

### Pitcher et al., Supplementary Figure 7

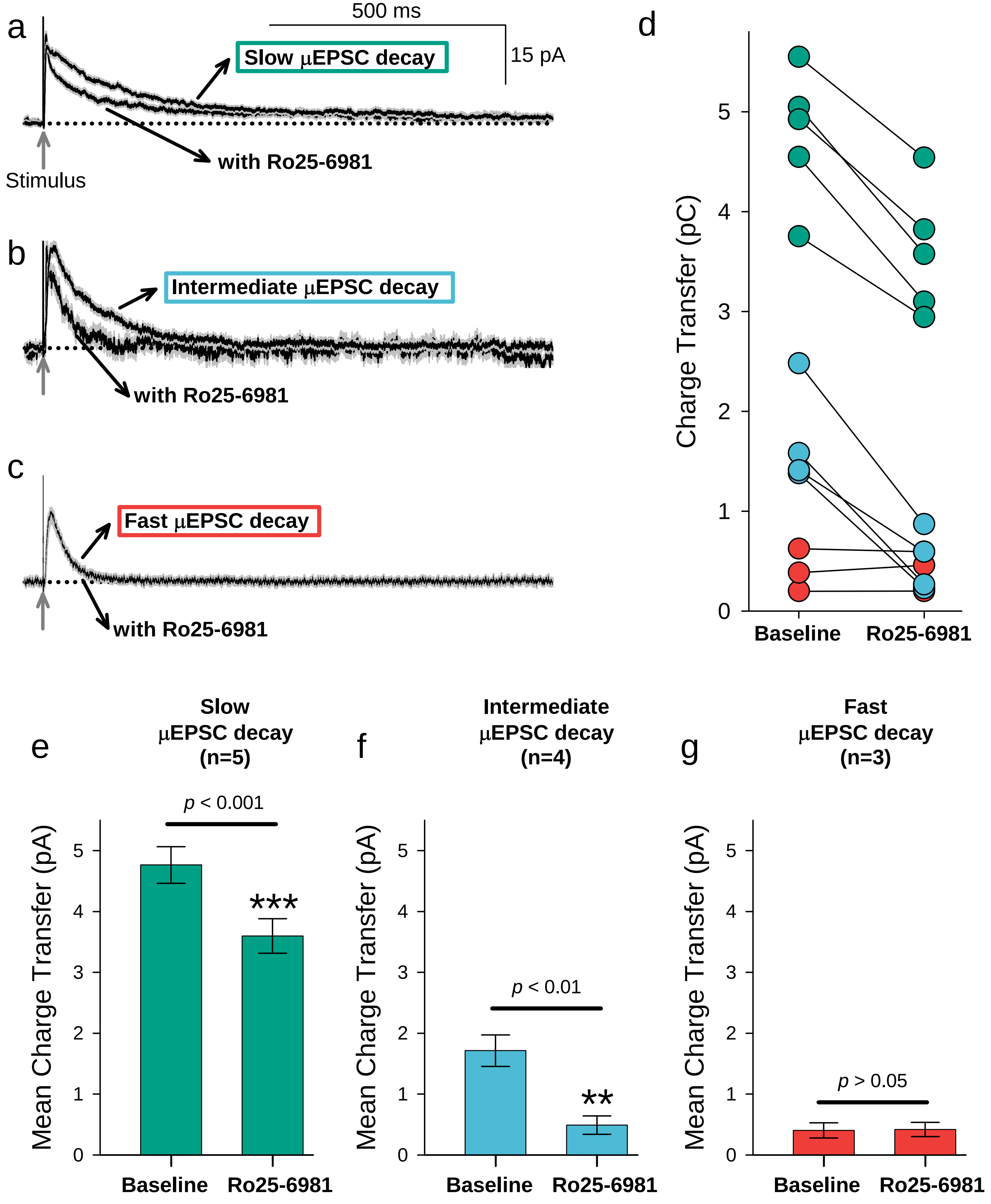

### Pitcher et al., Supplemenatry Figure 8

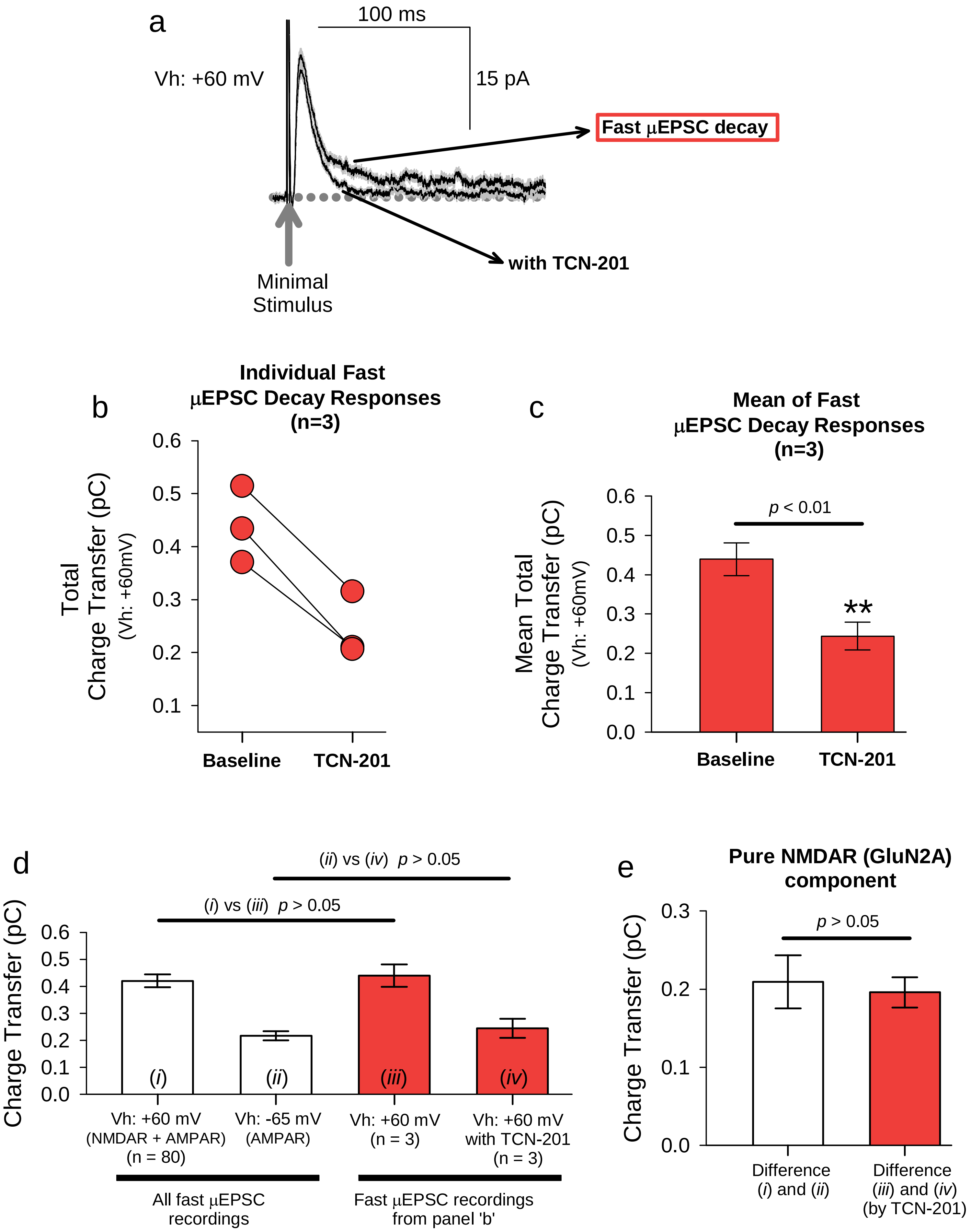

**Pitcher et al., Supplementary Figure 9**

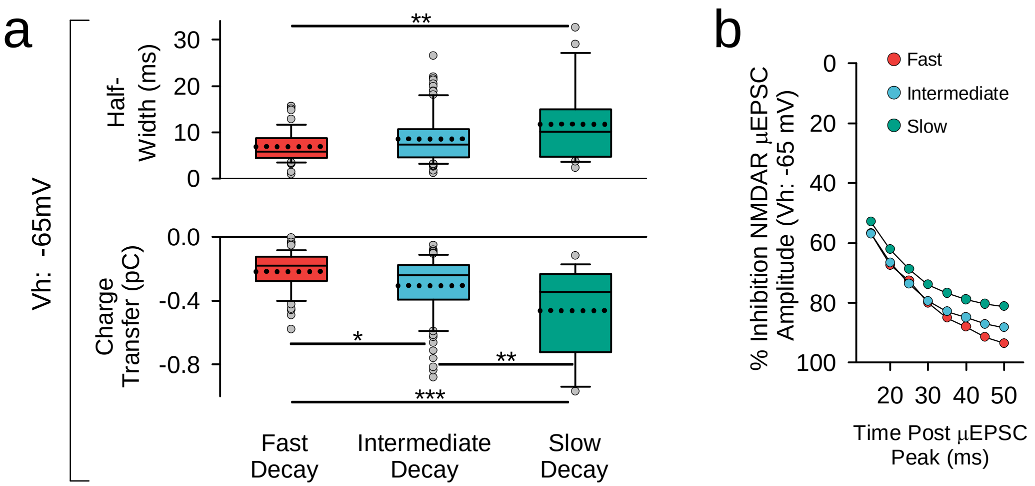

**Pitcher et al., Supplementary Figure 10**

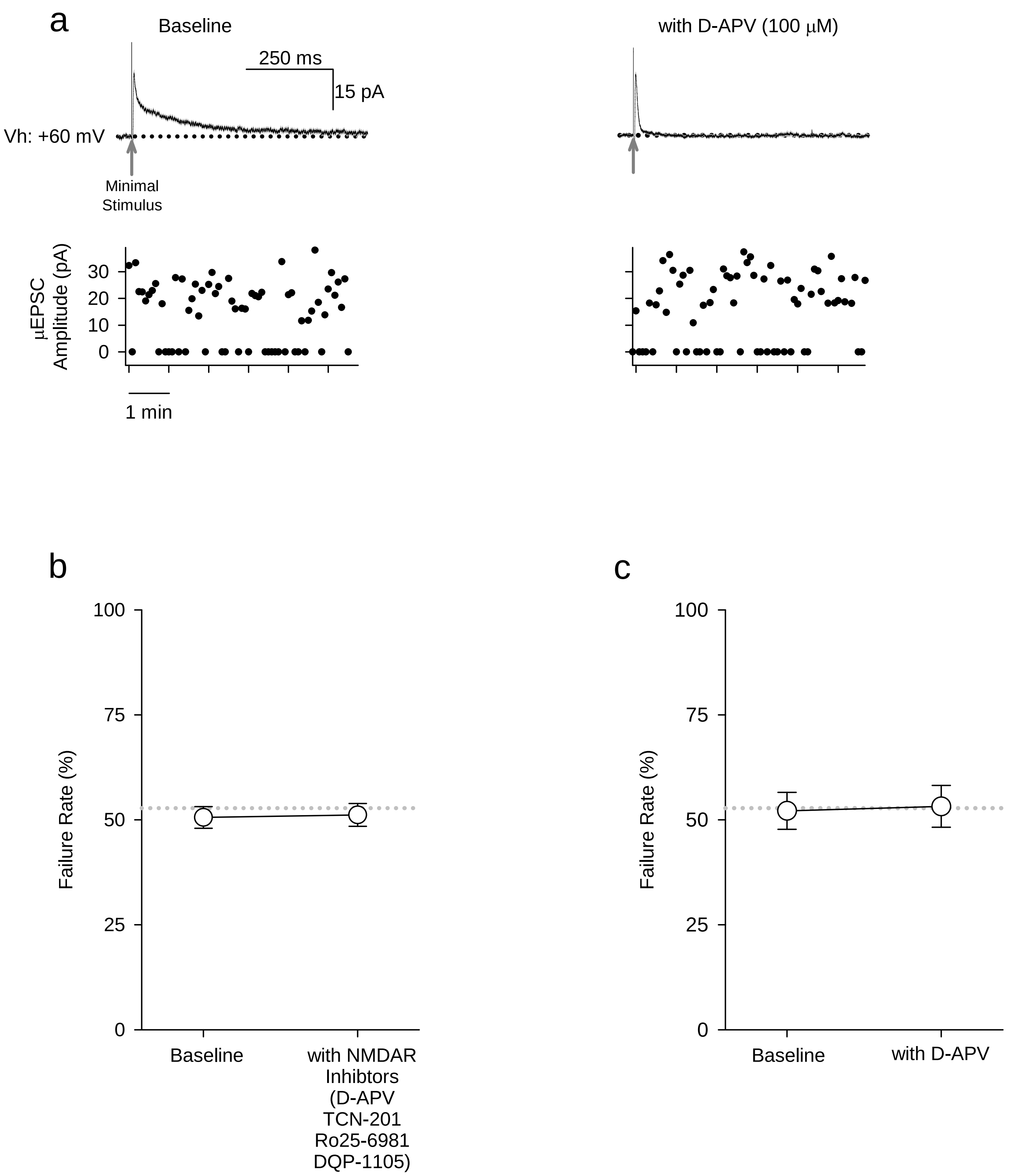

**Pitcher et al., Supplementary Figure 11**

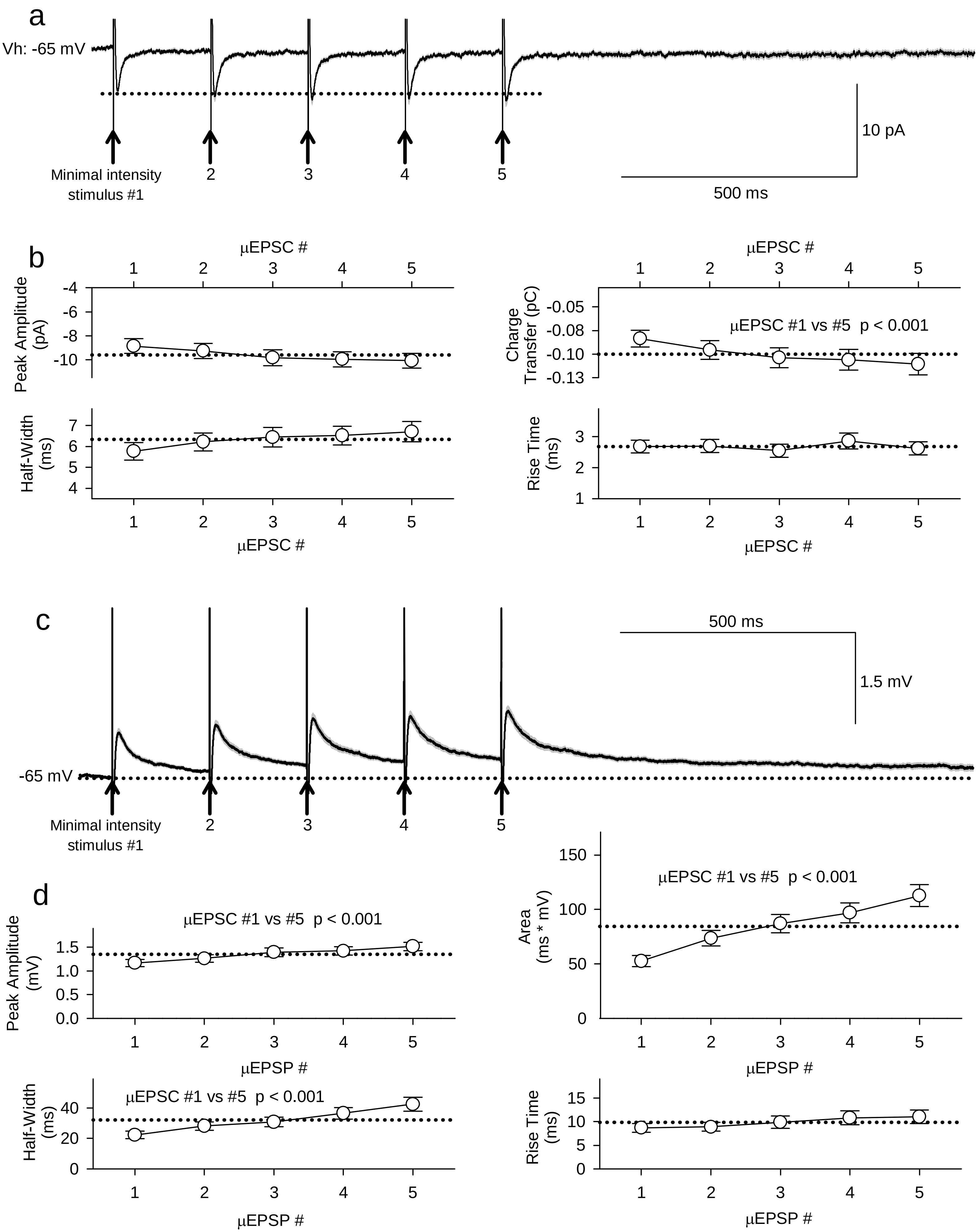

### Pitcher et al., Supplementary Figure 12

a

**Pure μEPSCs (successes only)**

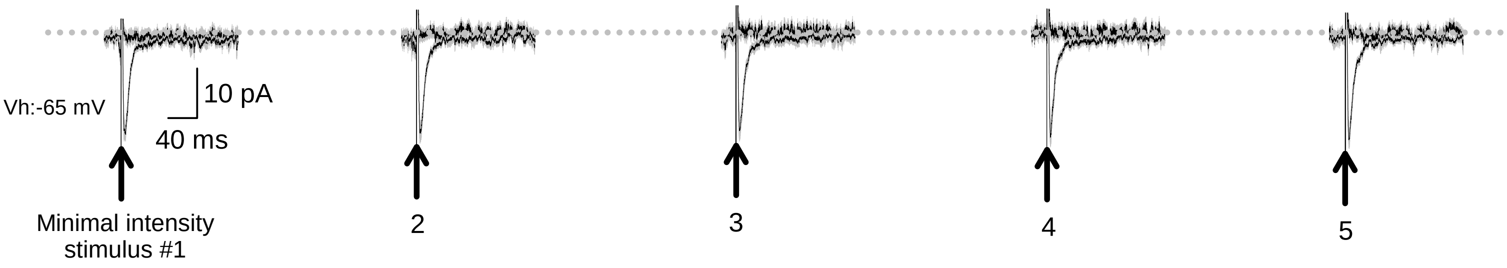


b

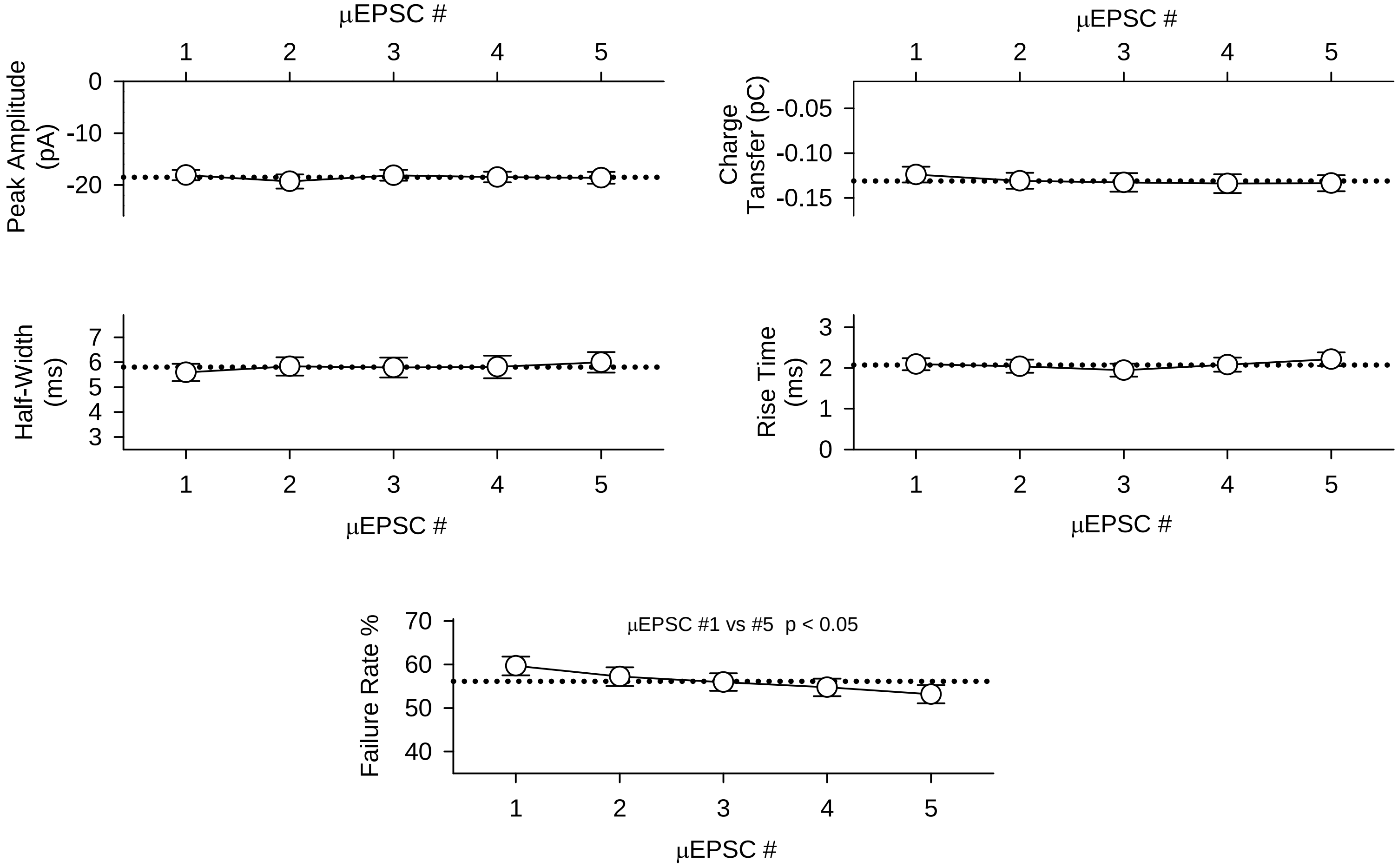

### Pitcher et al., Supplementary Figure 13

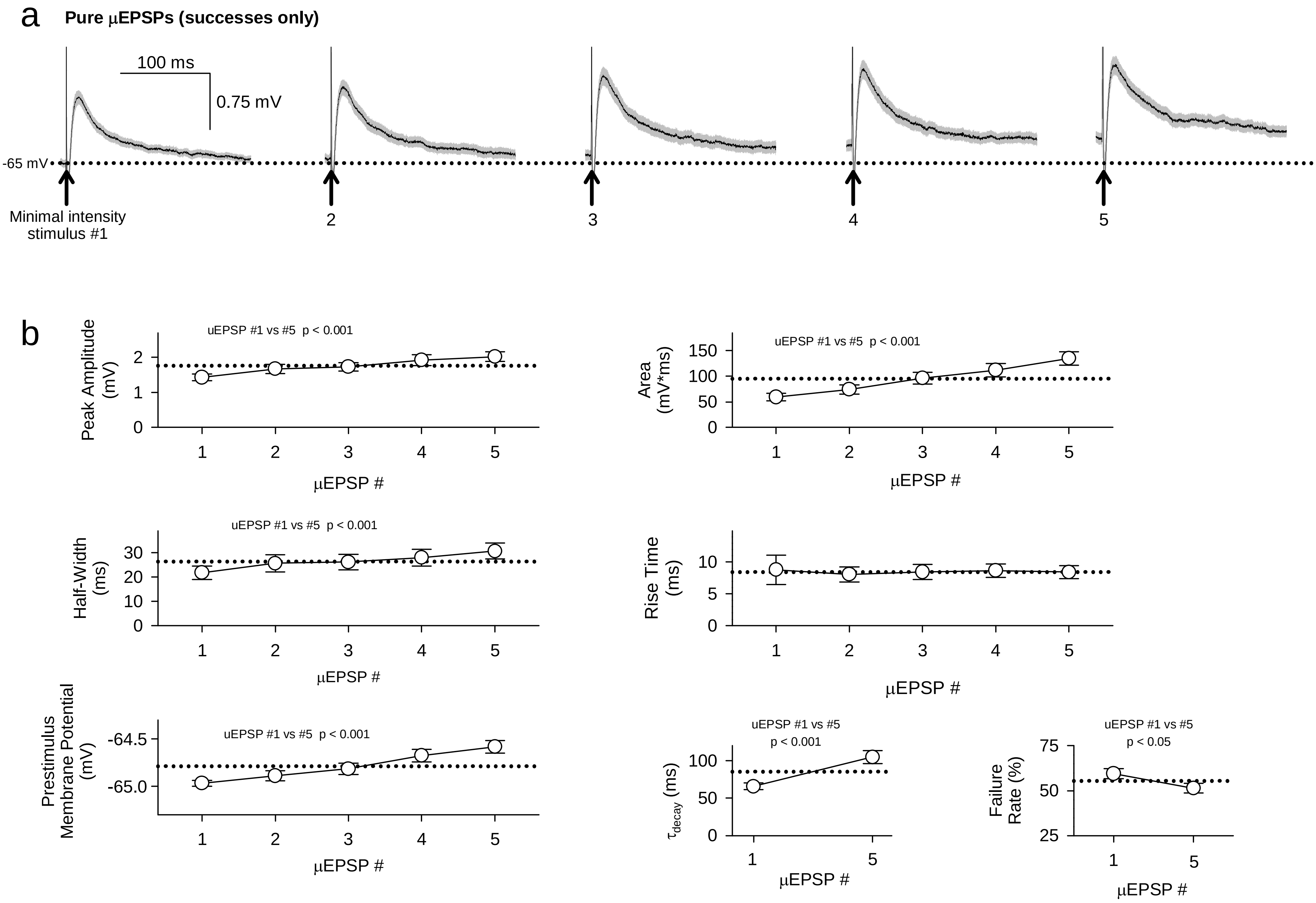

### Pitcher et al., Supplementary Figure 14

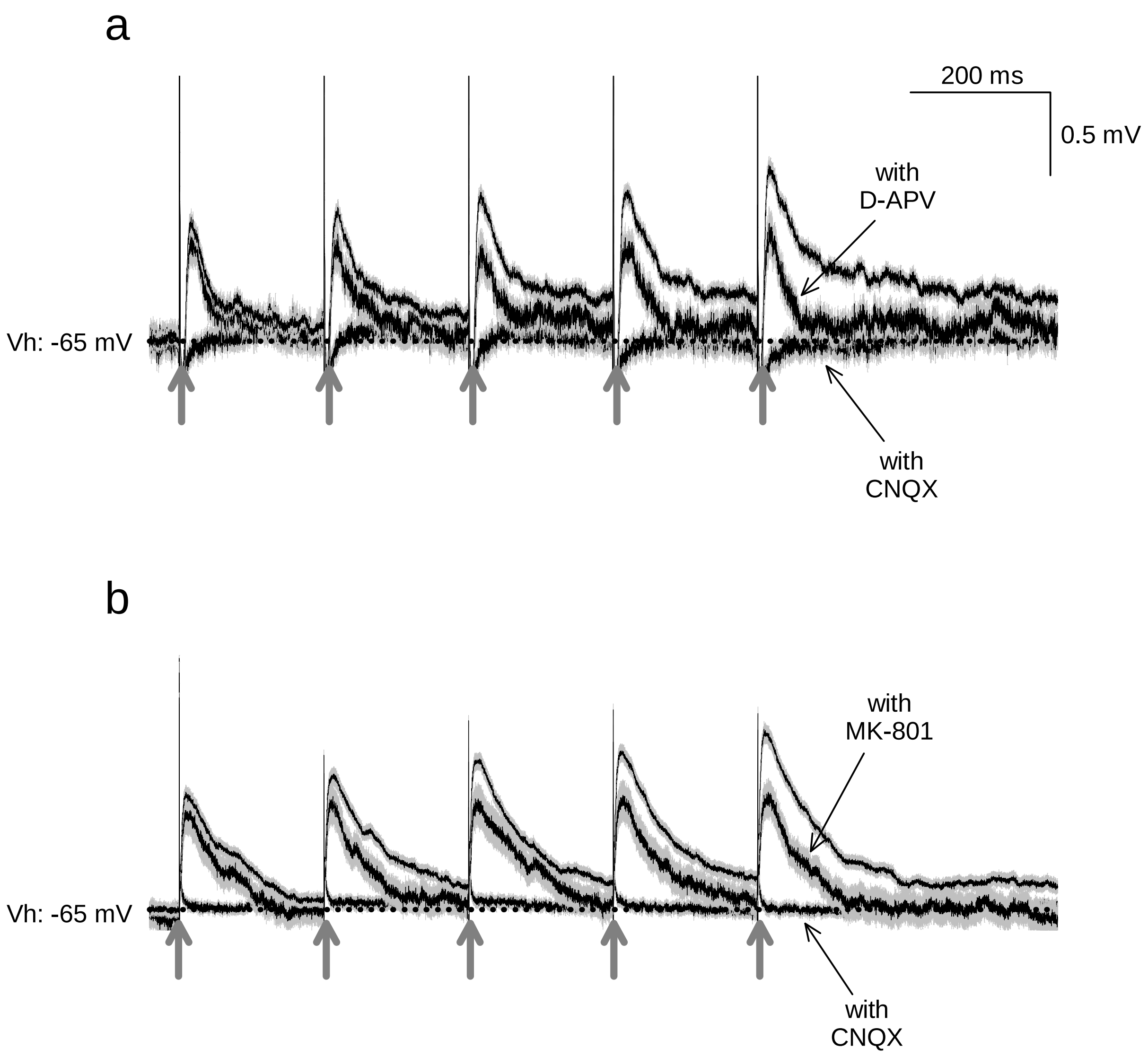
